# The Torsin/ NEP1R1-CTDNEP1/ Lipin axis regulates nuclear envelope lipid metabolism for nuclear pore complex insertion

**DOI:** 10.1101/2020.07.05.188599

**Authors:** Julie Jacquemyn, Joyce Foroozandeh, Katlijn Vints, Jef Swerts, Patrik Verstreken, Natalia V. Gounko, Sandra F. Gallego, Rose Goodchild

**Affiliations:** VIB-KU Leuven Center for Brain and Disease Research; Dept. of Neurosciences, KU Leuven, 3000 Leuven, Belgium; VIB-KU Leuven Center for Brain & Disease Research, Electron Microscopy Platform & VIB-Bioimaging Core, 3000 Leuven, Belgium

**Keywords:** AAA+ enzyme, dystonia, Lipin, TOR1A, TorsinA LAP1, TOR1AIP1, phosphatidic acid, diacylglycerol, triglyceride, lipid droplet, nuclear pore complex, nuclear membrane, nuclear envelope, endoplasmic reticulum, lipid synthesis, fat body, genetic suppressor, phosphatase, *Drosophila*

## Abstract

Torsin ATPases of the endoplasmic reticulum (ER) and nuclear envelope (NE) lumen inhibit Lipin-mediated phosphatidate (PA) to diacylglycerol (DAG) conversion by an unknown mechanism. This excess PA metabolism is implicated in *TOR1A*/TorsinA diseases, but it is unclear whether it explains why Torsin concomitantly affects nuclear structure, lipid droplets (LD), organelle and cell growth. Here a fly miniscreen identified that Torsins affect these events via the NEP1R1-CTDNEP1 phosphatase complex. Further, Torsin homo-oligomerization rather than ATPase activity was key to function. NEP1R1-CTDNEP1 activates Lipin by dephosphorylation. We show that Torsin prevents CTDNEP1 from accumulating in the NE and excludes Lipin from the nucleus. Moreover, this repression of nuclear PA metabolism is required for interphase nuclear pore biogenesis. We conclude that Torsin is an upstream regulator of the NEP1R1-CTDNEP1/ Lipin pathway. This connects the ER/NE lumen with PA metabolism, and affects numerous cellular events including it has a previously unrecognized role in nuclear pore biogenesis.

**Highlights:**

- Nuclear envelope PA-DAG-TAG synthesis is independently regulated by Torsin and Torip/LAP1
- Torsin removes CTDNEP1 from the nuclear envelope and excludes Lipin from the nucleus
- Excess nuclear envelope NEP1R1-CTDNEP1/ Lipin activity impairs multiple aspects of NPC biogenesis
- NEP1R1-CTDNEP1/ Lipin inhibition prevents cellular defects associated with *TOR1A* and *TOR1AIP1* / LAP1 disease

## Introduction

The endoplasmic reticulum (ER) is the largest organelle of the cell. It surrounds the nucleus as the inner and outer nuclear membranes (INM and ONM), and extends throughout the cell body. Most cellular lipids are synthesized by enzymes localized on or in the ER membranes. However, it is unclear how lipid synthesis is spatially organized within the ER, including that several lipid enzymes localize in the nucleus and/ or the INM (Jacquemyn et al., 2017).

Lipin dephosphorylation of PA into DAG is a key node in lipid synthesis. The DAG production drives triglyceride (TAG) and lipid droplet (LD) production (Adeyo et al., 2011), while PA removal suppresses membrane lipid synthesis (Craddock et al., 2015). In this manner, Lipin tends to promote energy storage at the expense of organelle and cell growth. Numerous stimuli modify lipid metabolism through Lipin. This includes the cell cycle (Bahmanyar et al., 2014), mechanical stress (Romani et al., 2019), nutrition (Peterson et al., 2011), anoxia (MacVicar et al., 2019), membrane composition (Eaton et al., 2013; Karanasios et al., 2010), and when cells alter morphology (Yang et al., 2020). Lipin is a large > 100kD soluble enzyme (Khayyo et al., 2020) that receives extensive post-translational modifications: acetylation, sumoylation, ubiquitination, and phosphorylation have all been described (Harris et al., 2007; Li et al., 2018; Liu and Gerace, 2009; O’Hara et al., 2006; Shimizu et al., 2017). Lipin PA phosphatase activity requires that the N-terminus inserts into the membrane as an amphipathic helix (Khayyo et al., 2020). This event is inhibited by phosphorylation, which reduces Lipin PA phosphatase activity (Eaton et al., 2013; Huffman et al., 2002; Karanasios et al., 2010; O’Hara et al., 2006). Lipin phosphorylation also counteracts a nuclear localization signal and localizes the enzyme in the cytosol rather than nucleus (Peterson et al., 2011). Several kinases, most notably mTORC1, regulate Lipin through phosphorylation state (Hennessy et al., 2019; Huffman et al., 2002; Peterson et al., 2011). They are opposed by phosphatases (Kok et al., 2014; Okuno et al., 2019), including an unusual transmembrane phosphatase (**C-T**erminal **D**omain **N**uclear **E**nvelope **P**hosphatase (CTDNEP1) and its regulatory subunit (NEP1R1)) (Han et al., 2012; Kim et al., 2007; Santos-Rosa et al., 2005).

Torsin AAA+ ATPases of the ER/ NE lumen are newly uncovered Lipin regulators. This was identified in the *Drosophila* fat body which performs functions similar to liver and adipose tissue (Grillet et al., 2016). Torsins are highly conserved: all domains and key residues are conserved between worm, fly, and mammalian TorsinA and TorsinB (Sosa et al., 2014). TorsinA is also a disease protein. Homozygous *TOR1A*/TorsinA mutations cause a recessive neurological syndrome (Kariminejad et al., 2017), while a heterozygous *TOR1A*/TorsinA mutation causes the neurological movement disorder of dystonia (Ozelius et al., 1997). It is also clear that mammalian TorsinA negatively regulates Lipin, including there is excess Lipin activity in patient cells (Cascalho et al., 2020).

The mechanism connecting Torsins with Lipin remains unclear, and they reside on opposite faces of the ER membrane. Torsins form at least two different complexes inside the ER. They interact with transmembrane “Torsin Activating Proteins” that stimulate their ATPase activity, and may also transduce information across the membrane (Fig 1A). In mammals these are LAP1 of the INM (*TOR1AIP1*), and LULL1 of the main ER (*TOR1AIP2*) (Chase et al., 2017; Sosa et al., 2014; Zhao et al., 2013). LAP1 loss causes a lethal multisystemic congenital syndrome (Fichtman et al., 2019), again highlighting that this evolutionarily conserved protein network has clinical as well as biological significance. Alternatively, Torsin monomers assemble into ATPase inactive “split ring”-like structures (Chase et al., 2017). These are sufficiently stable *in vitro* that they recruit additional Torsin monomers and extend into helical polymers (Demircioglu et al., 2019). Their functional importance remains unclear.

**Figure 1.**
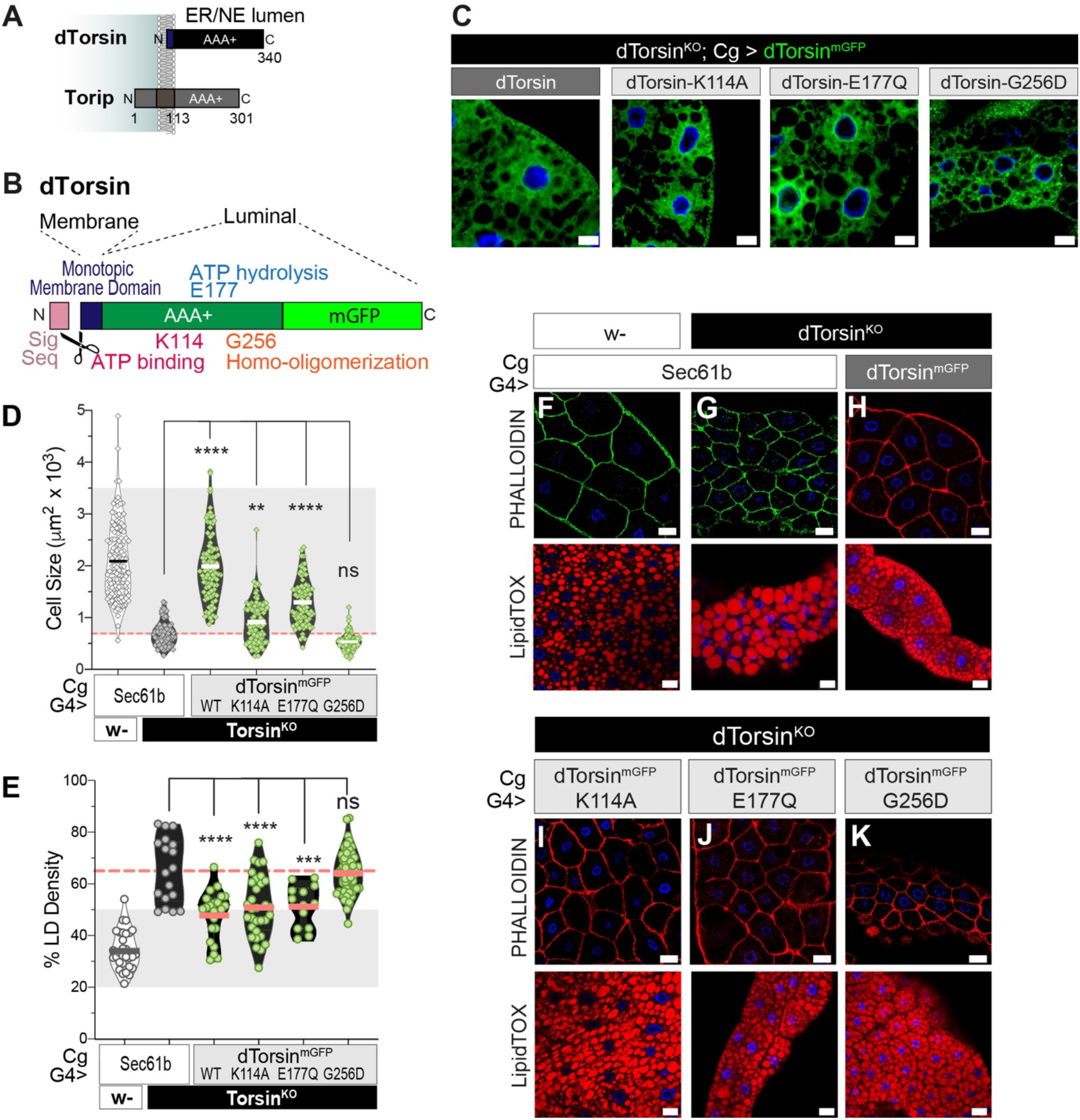
dTorsin homo-oligomerization is required for fat body cell growth and LD homeostasis. A & B) dTorsin and Torip topology, fusion, and mutants. C) Wild-type and mutant dTorsin^mGFP^ proteins expressed in the *dTorsin^KO^* fat body. Scale bars, 10μm. D & E) Points show values from individual fat body cells, bars show the group mean. Light grey highlights 2x SD of the control mean, and red lines show *dTorsin^KO^* mean. F – K) Phalloidin and LipidTOX staining reveal the cell periphery and LD, respectively, in fat body cells expressing Sec61, dTorsin, or dTorsin mutants. Scale bars, 20μm.

Torsins impact numerous cellular events and it is unclear whether this reflects multiple functions, or a core function with many downstream consequences. This includes that Torsins and LAP1 are best known for roles in the nuclear envelope (NE). Torsin loss impairs nuclear pore complex (NPC) biogenesis and levels in worm and mammals (Rampello et al., 2020; VanGompel et al., 2015). Torsin and LAP1 loss also produce abnormal vesicle-like membranes in the NE lumen (Goodchild et al., 2005; Kim et al., 2010) which potentially represent stalled nuclear egress vesicles (Jokhi et al., 2013), or intermediates of NPC insertion (Rampello et al., 2020). These phenomena underlie the theory that Torsin-LAP1 ATPase activity directly induces membrane scission (Jokhi et al., 2013). However, this has not been reconciled with the knowledge that Torsin inhibits Lipin, or that TorsinA and LAP1 both suppress TAG deposition in mouse liver (Shin et al., 2019). In fact, PA-DAG-TAG synthesis takes place at the yeast INM and, although this has no known role for the NPC or egress vesicles, it can affect nuclear membrane structure (Romanauska and Kohler, 2018).

Here we make the surprising finding that Torsin homo-oligomers regulate lipid metabolism. Torsin does not require the LAP1 (Torip) ATPase activator of fly, and LAP1/Torip regulates lipid metabolism without requiring Torsin. We further identify that Torsin negatively regulates the NE/ nuclear levels of CTDNEP1 and Lipin, respectively, and that the NEP1R1-CTDNEP1 phosphatase complex indeed underlies how Torsin regulates LD, cell growth, organelle size, and nuclear membrane structure. This includes that excess NEP1R1-CTDNEP1/ Lipin activity prevents at least two different steps of NPC biogenesis. We conclude that Torsin is part of the NEP1R1-CTDNEP1/ Lipin axis and, as a pathway, this couples the ER/NE lumen to lipid metabolism for a remarkably broad set of cellular events. This includes a previously unrecognized role in NPC biogenesis, as well as explaining multiple *TOR1A* and *TOR1AIP1* disease-associated phenomenon.

## Results

### dTorsin homo-oligomers and Torip/LAP1 independently regulate lipid metabolism

*dTorsin* is on the X-chromosome and we refer to *dTorsin^-/y^* males as *dTorsin^KO^*. Late stage (5 days-old; 5DO) third instar *dTorsin^KO^* larvae have an abnormally small fat body, concomitant with poor cell growth and over production of TAG and LD (Grillet et al., 2016). We explored how dTorsin regulates fat body lipid metabolism using a series of mutants (Fig. 1B) with impaired ATP binding (Walker A K114A), ATP hydrolysis (Walker B E177Q), or homo-oligomerization (back interface G256D that retains binding to Torsin Activators (Chase et al., 2017)). The Cg-Gal4 driver expressed similar levels of each dTorsin^mGFP^ in the in the ER/NE of *dTorsin^KO^* fat body cells (Fig. 1C). As expected, dTorsin^mGFP^-WT expression returned *dTorsin^KO^* cells to the same size as control animals, and strongly suppressed the LD defect (Fig. 1D & E, Fig. 1F - H). dTorsin^mGFP^-K114A and dTorsin^mGFP^-E177Q also rescued *dTorsin^KO^* cell size compared to *dTorsin^KO^* expressing the control Sec61b ER protein (Fig. 1D, 1I & J), and were as effective as dTorsin^mGFP^-WT at reducing the overly dense LD in *dTorsin^KO^* cells (Fig. 1E, 1I & J). Thus, mutants with impaired nucleotide binding and hydrolysis retained activity. In contrast, dTorsin^mGFP^-G256D failed to improve *dTorsin^KO^* cell size (Fig. 1D & 1K) or LD density (Fig. 1E & 1K). Surprisingly, this points to Torsin homooligomers as the structure that regulates lipid metabolism.

dTorsin, like all Torsins, lacks the catalytic arginine residue needed for ATP hydrolysis, which is instead present in their Activator partner (Sosa et al., 2014). Consequently, ATPase activity only occurs when Torsin binds an Activator. There is one Activator in *Drosophila*, Torip (Fig. 1A; (Sosa et al., 2014)) which resides in the NE, analogous to mammalian LAP1 (Fig. 2A; (Grillet et al., 2016)). We first addressed whether Torip is the NE binding partner of dTorsin by introducing an 11 base-pair deletion into *Torip* exon1. This removed *Torip* mRNA, and introduced a premature stop codon before the transmembrane domain (Fig. S1A-C). dTorsin^mGFP^-E177Q concentrated at the NE of wild-type cells, which is typical since ATP-bound Torsins strongly interact with their INM Activator (Fig. 2B; (Naismith et al., 2009)). In contrast, dTorsin^mGFP^-E177Q was no longer localized at the NE when Torip levels were reduced by *Torip*^Δ11/+^ (Fig. 2C). We therefore conclude that *Torip*^Δ11/+^ is a null allele, and Torip is the LAP1 ortholog of fly. From here on we refer to *Torip*^Δ11/Δ11^ as *Torip^-/-^*.

**Figure 2.**
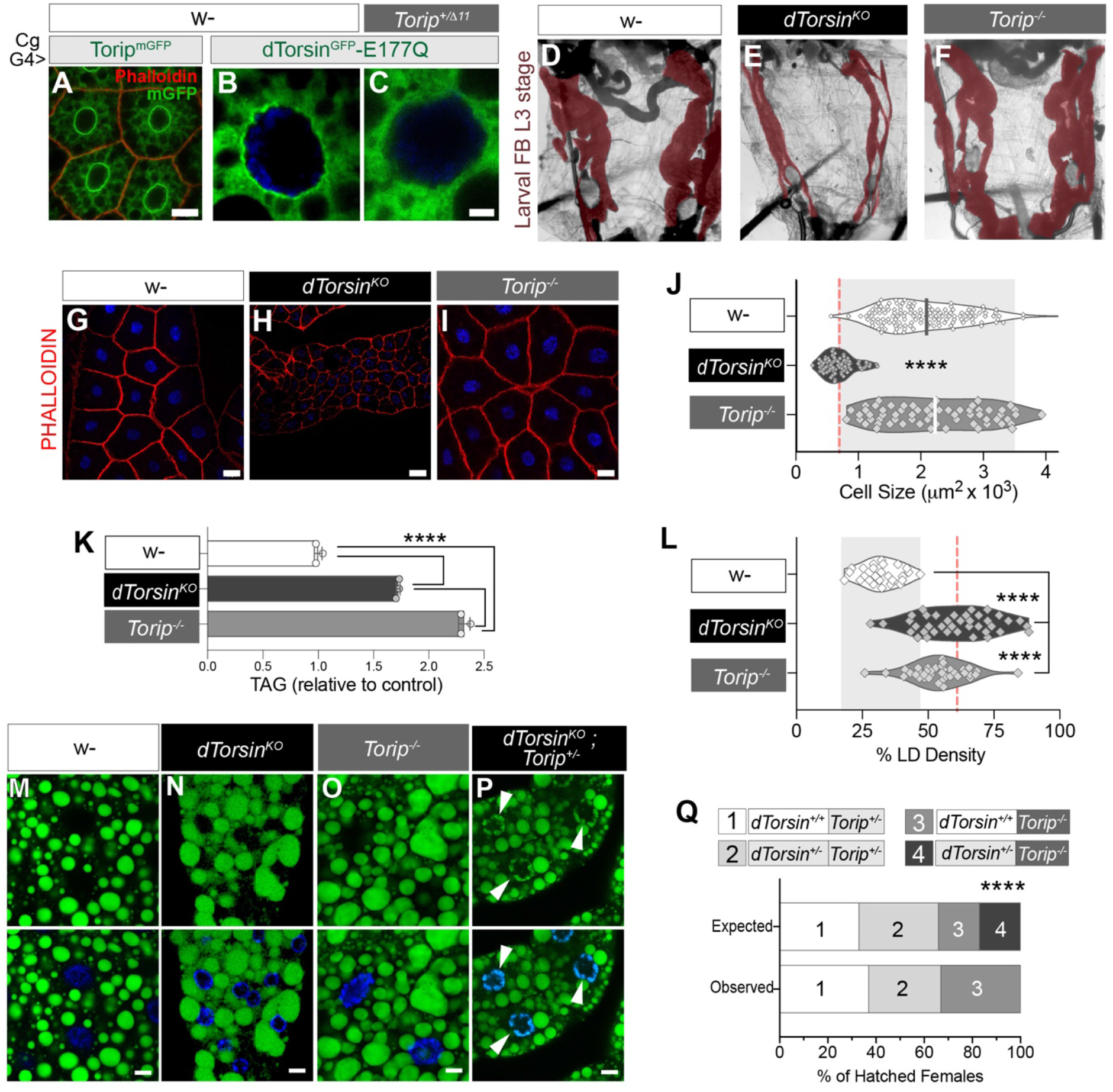
*dTorsin* and *Torip/LAP1* independently regulate lipid metabolism at the nuclear envelope. A - C) Torip^mGFP^ or dTorsin^mGFP^ in the wild-type (w-) or *Torip*^+/Δ11^ fat body of 3^rd^ instar fly larvae. Scale bars, 20μm and 5μm respectively. D – F) Control, *dTorsin^KO^*, and *Torip^-/-^* larvae with pseudo-colored fat body. G – I) Phalloidin staining reveals the cell periphery of control, *dTorsin^KO^*, and *Torip^-/-^* fat body cells. Scale bars, 20 μm. J) Points show the area of individual fat body cells and bars show the mean. Light grey is 2x SD of the control mean, red line shows *dTorsin^KO^* mean. K) Mean ± SD of TAG present in 3DO larvae. TAG was measured in three sets of thirty larvae. L) The percentage of a 5DO fat body cell labeled by BODIPY. Points show density in individual cells, and bars show mean. Light grey is 2x SD of the control mean, red line shows *dTorsin^KO^* mean. M - P) Neutral lipid staining of the 5DO larval fat body. Scale bars, 10 μm. Q) % of females that hatch from mating *dTorsin^+/-^*;; *Torip^-^*/TM6C females with y/FM7i;; *Torip^-^*/TM6C males.

*dTorsin^KO^* larvae have a smaller fat body than wild-types (Fig. 2D vs E). In contrast, *Torip^-/-^* fat body size was normal (Fig. 2F). In addition, and unlike the *dTorsin^KO^*, there was no difference between the size of *Torip^-/-^* fat body cells and those of wild-type (Fig. 2G - J). This shows that *dTorsin* retains activity in the absence of *Torip*, consistent with the importance of the G256 back-interface residue. However, the normal appearance of *Torip^-/-^* animals is surprising since LAP1 suppresses TAG deposition in the mouse liver (Shin et al., 2019). We therefore specifically examined TAG in *dTorsin^KO^* and *Torip^-/-^* larvae. Both had more TAG than controls (Fig. 2K), and neutral lipid staining detected that their cells were overly packed with LD compared to controls (Fig. 2L - O). Thus, *dTorsin* and *Torip* indeed both suppress TAG production like their mammalian counterparts. However, it appears that they operate independently given the divergence of the two null phenotypes. We tested this by intercrossing the alleles. We failed to generate *dTorsin^KO^;;Torip^-/-^* larvae, although did identify L3 stage *dTorsin^KO^*;;*Torip^+/-^*. These had small fat body cells like the *dTorsin^KO^*, but also gained the additional defect of nuclear localized neutral lipid staining (Fig 2P; arrows). Moreover, *Torip* loss was incompatible with *dTorsin^+/-^* development (Fig 2Q). We therefore conclude that both *dTorsin* and *Torip/LAP1* inhibit TAG synthesis, but they do so independently. Moreover, the fact that combined loss produces nuclear TAG shows that they both perform this function at the NE.

### dTorsin regulates lipid metabolism through the transmembrane NEP1R1-CTDNEP1 phosphatase complex

We previously showed that *Lipin* RNAi partially restored *dTorsin^KO^* fat body size (Grillet et al., 2016). To identify other components of the *dTorsin* pathway we performed an RNAi survey of transmembrane lipid enzymes, regulators of lipid metabolism, and previously described Torsin interaction partners that might convey information on Torsin structure across the membrane. We expressed individual UAS-RNAi lines using r4-Gal4 that is highly active in the fat body (Lee and Park, 2004). Most lines had no impact on the 5DO *dTorsin^KO^*, including RNAi against the *Koi* Sun protein (Fig 3A). In contrast, RNAi against (1) the *Ctdnep1* protein phosphatase, (*Dullard, CG1696*), and (2) its regulatory subunit (*Nep1r1; CG41106*) dramatically improved *dTorsin^KO^* fat body size (Fig 3A). In fact, *Ctdnep1* and *Nep1r1* RNAi appeared similarly effective as dTorsin-WT re-expression (Fig. 3A; final panel). We further confirmed that *Ctdnep1* and *Nep1r1* RNAi rescued *dTorsin^KO^* cell size, as did *Lipin1* RNAi (Fig. 3B, Fig. S2A), while none affected cell size in wild-type animals (Fig. S2B).

**Figure 3.**
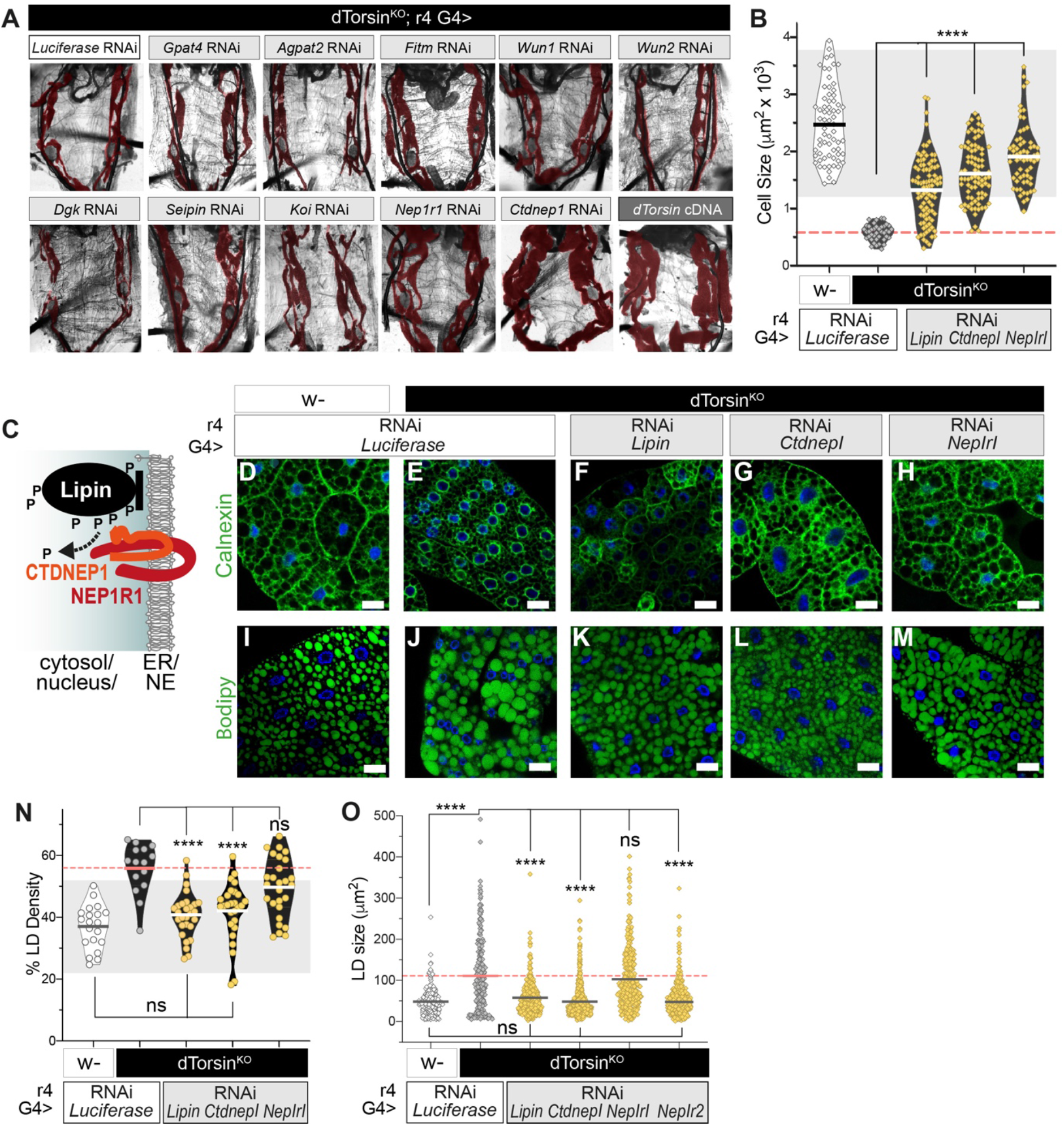
*Nepr1/2-Ctdnep1* RNAi suppresses cell growth, organelle volume, and LD defects of *dTorsin^KO^* larvae. A) *dTorsin^KO^* expressing RNAi against candidate genes implicated in Torsin or Lipin function, or expressing wild-type *dTorsin* (control). The fat body is pseudo-colored red. B) Points show size of individual fat body cells, bars show the mean. Light grey shading highlights 2x SD of the control mean, and red line shows *dTorsin^KO^* mean. C) Topology of CTDNEP1-NEP1R1 and Lipin. D - M) Fat body cells labeled by anti-Calnexin or BODIPY detection of neutral lipids. Scale bars, 20μm. N) Points show values from individual fat body cells, bars show group mean. Light grey highlights 2x SD of the control mean, and dotted red lines show the *dTorsin^KO^* mean. O) Points show the size of individual LD, bars show group means, and red line shows the *dTorsin^KO^* mean. Group differences were detected with the non-parametric Kruskal-Wallis test.

NEP1R1 regulates CTDNEP1 protein phosphatase activity that, in turn, regulates Lipin PA phosphatase activity (Fig. 3C) (Han et al., 2012; Kim et al., 2007; Su et al., 2014). *Lipin, Ctdnep1* and *Nep1r1* RNAi similarly reduced their mRNAs (Fig. S2C). Thus, the fact that *Nep1r1* and *Ctdnep1* RNAi prevent *dTorsin^KO^* defects as effectively as *Lipin* RNAi, very strongly implicates NEP1R1-CTDNEP1 gain-of-function as the mechanism connecting Torsin to Lipin.

We further explored whether *Nep1r1/ Ctdnep1/ Lipin* pathway gain-of-function explains why *dTorsin* affects lipids. We previously showed that abnormally low membrane lipid levels in the *dTorsin^KO^* occur in parallel with a less dense ER (Grillet et al., 2016). We detected ER density with anti-Calnexin. While ER extended throughout wild-type cells, anti-Calnexin primarily labeled the perinuclear region in *dTorsin^KO^* cells and there was minimal signal in the cell periphery (Fig. 3D vs E). *Lipin, Ctdnep1*, and *Nep1r1* RNAi all normalized *dTorsin^KO^* Calnexin staining towards a wild-type distribution (Fig. 3F – H), suggesting pathway gain-of-function indeed underlies the ER defects of the *dTorsin^KO^*. Next, we examined *dTorsin^KO^* TAG overproduction that produces larger and more densely packed LD (Fig. 3I – J). The three RNAi transgenes returned neutral lipid staining of *dTorsin^KO^* cells towards a wild-type state (Fig. 3K – M). Quantitation showed that *Lipin* and *Ctdnep1* RNAi reduced *dTorsin^KO^* LD density and size to wild-type levels (Fig. 3N & O). Surprisingly, *Nep1r1* RNAi did not. This suggested that the second *Nep1r* gene of *Drosophila* (CG8009; *Nep1r2*) may also affect LD. Indeed, *Nep1r2* RNAi normalized *dTorsin^KO^* LD size to wild-type levels (Fig. 3O).

### *Nep1r1-Ctdnep1* gain-of-function explains why *dTorsin* is required for NPC biogenesis

Torsin activity is important at the NE. Now, the fact we identified a) *dTorsin* suppresses NE-localized TAG production, and b) appears to regulate lipid metabolism via the NE/ER localized NEP1R1-CTDNEP1 complex, led us to explore whether the same pathway was responsible for the NE importance of Torsins. We focused on the role of Torsin in the poorly understood process of interphase NPC insertion (Fig. 4A & B; (Rampello et al., 2020)). Notably, fat body cells exit the cell cycle after embryogenesis (Zheng et al., 2016) and thus, are ideal to study interphase events without interference from mitotic NE breakdown.

**Figure 4:**
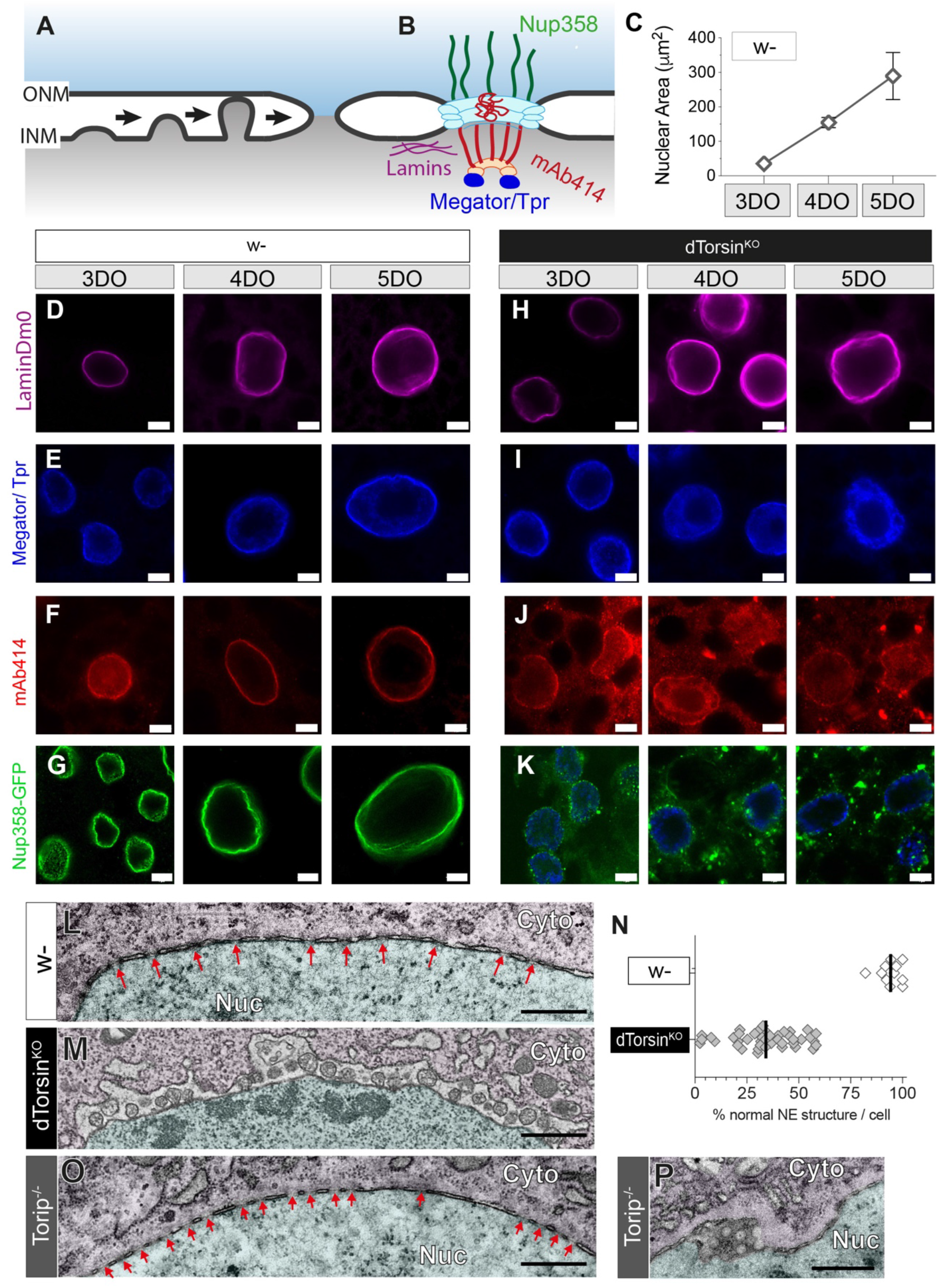
NPC insertion into the nuclear membranes requires *dTorsin* but not *Torip*. A) Interphase NPC insertion occurs via inside-out extrusion of the INM, followed by INM / ONM fusion (Otsuka et al., 2016). Nucleoporins of immature pores are not depicted. B) The NPC, highlighting proteins examined in fat body cells. C) Fat body nuclei increase in size over three days of 3^rd^ instar development. D – K) Fat body cell nuclei in control w-(D – G) or *dTorsin^KO^* (H – K) during the 3^rd^ instar larval period, and labeled with anti-LaminDm0, Megator/Tpr, mAb414, or expressing a Nup358-GFP transgene with Cg-Gal4. Scale bars, 5μm. L & M) Representative TEM of the NE of control and *dTorsin^KO^* 5DO fat body cells. Green highlights nucleus, magenta highlights cytosol, red arrows show correctly inserted NPC. Nuc, nucleus; Cyto, cytosol. Scale bars, 500nm. N) The percentage of NE in individual 5DO fat body cells with normal INM, ONM, and NPC morphology. Bars show group mean. O & P) TEM of NE structure in 5DO *Torip^-/-^* fat body cells.

Fat body nuclei expanded over development (Fig. 4C & D) and concomitantly maintained levels of the Megator/Tpr NPC basket component (Fig. 4E), FG-repeat channel Nups detected by mAb414 (Fig. 4F), and cytoplasmic Nup358 (Fig. 4G) (Weberruss and Antonin, 2016). Thus, fat body nuclei insert NPC during interphase growth. *dTorsin^KO^* fat body nuclei also expanded over development, and maintained an intact nuclear lamina (Fig. 4H). However, while Megator/Tpr localized to the *dTorsin^KO^* NE in 3DO and 4DO larvae, it was primarily in the nuclear interior at 5DO (Fig. 4I). mAb414 barely labelled the *dTorsin^KO^* NE at the three ages (Fig. 4J), and there was far less Nup358-GFP in the NE of *dTorsin^KO^* cells than controls (Fig. 4G vs. K). We assessed NE ultrastructure. Wild-type cells had the normal parallel arrangement of INM and ONM with intermittent NPC (Fig. 4L). In contrast, nuclei of 5DO *dTorsin^KO^* cells were coated with the classic membrane herniations of Torsin loss (Fig. 4M), to where < 30% of the NE was intact (Fig. 4N). Similar herniations contain nucleoporins in mammals (Rampello et al., 2020), and are reminiscent of immature NPC (Otsuka et al., 2016). We also examined *Torip^-/-^* animals. The majority of *Torip^-/-^* nuclei were normal and none had the INM herniations of Torsin loss (Fig. 4O). However, there were unusual circular profiles in place of well-defined INM and ONM in some *Torip^-/-^* cells (Fig. 4P). Thus, we conclude that *dTorsin* is required for NPC biogenesis in interphase fat body cells. In contrast, Torip loss only mildly affects the NE, including that NPC insertion appears relatively normal.

We examined if *Nep1r1/ Ctdnep1* gain-of-function underlies why Torsin loss prevents interphase NPC insertion. *dTorsin* re-expression restored NPC density to the 5DO *dTorsin^KO^* and removed all membrane defects (Fig. 5A & B, white vs black bars; Fig. 5C vs D), showing it is possible to genetically restore NE structure. *Ctdnep1* RNAi also strongly improved *dTorsin^KO^* NPC density and nuclear structure compared to animals expressing a control RNAi (Fig. 5A & B, Fig. 5E). Even more strikingly, *Nep1r1* RNAi restored NPC density to the same degree as *dTorsin* re-expression (Fig. 5A & F), including it almost completely removed nuclear membrane defects (Fig. 5B & F). We therefore conclude that Nep1r1 and Ctdnep1 gain-of-function causes the abnormal NPC biogenesis of cells lacking dTorsin.

**Figure 5.**
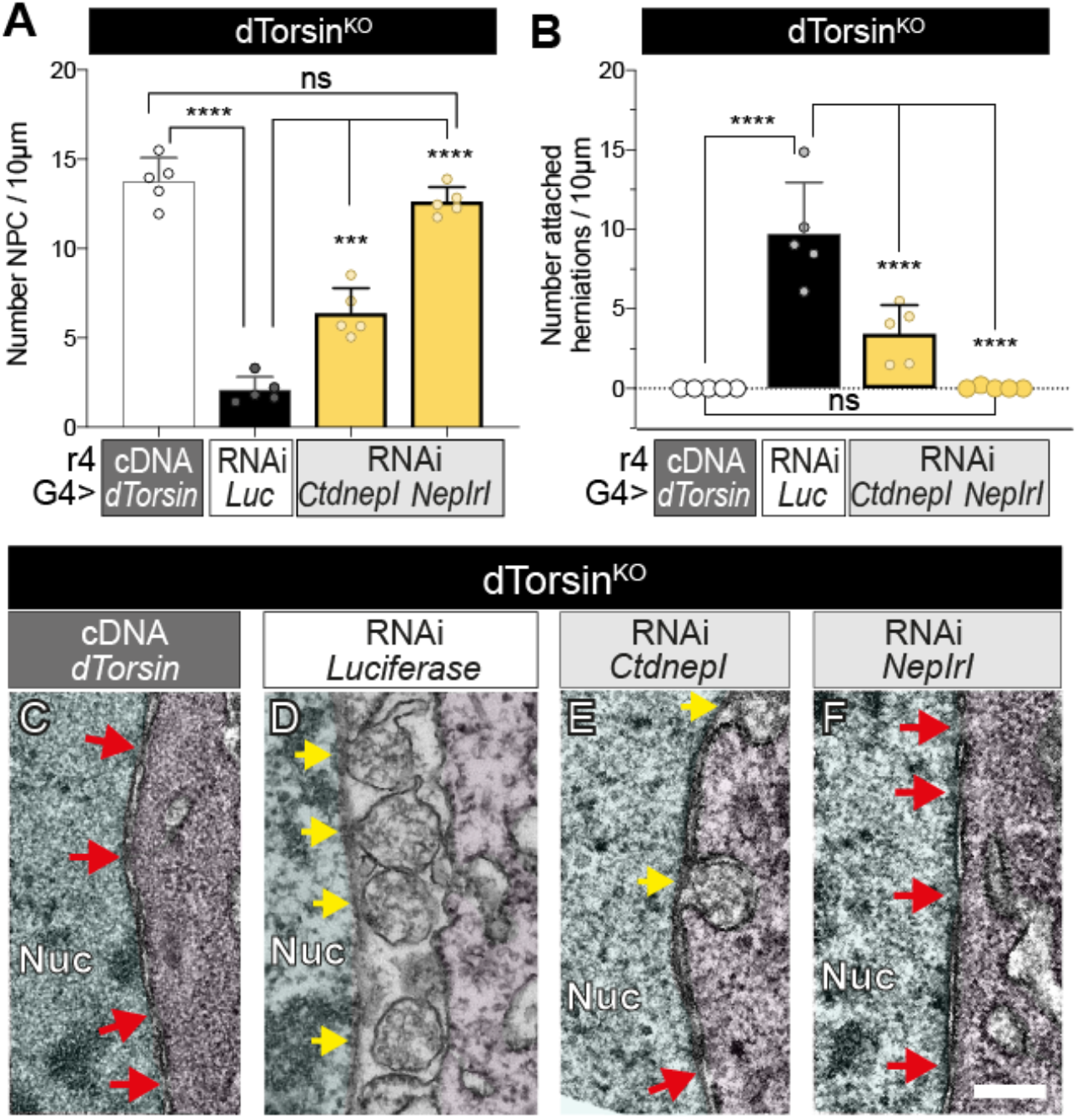
CTDNEP1-NEP1R1 RNAi restore interphase NPC insertion and suppress membrane defects in the *dTorsin^KO^* NE. A & B) Mean and SD of NPC and herniation density in the 5DO *dTorsin^KO^* NE. Individual points are the mean of multiple measurements from individual nuclei. C – F) NE ultrastructure in 5DO *dTorsin^KO^*. Red arrows, NPC, Yellow arrows, abnormal NE herniations. Scale bar, 500nm.

### Torsins control the nuclear localization of CTDNEP1 and Lipin

We sought the molecular basis of *Nep1r1* and *Ctdnep1* gain-of-function in *dTorsin^KO^* larvae. mRNA levels were no different between control and *dTorsin^KO^* animals (Fig. S3A - C). We therefore considered whether Torsin affected levels or localization by expressing the two proteins fused to GFP in the fly fat body. NEP1R1^mGFP^ primarily colocalized with Calnexin in control and *dTorsin^KO^* fat body cells, although some signal appeared to concentrate at the *dTorsin^KO^* NE. This NE concentration was specific to the *dTorsin^KO^*, and appeared to exceed NE-localized Calnexin signal (Fig. 6A & B; arrows) suggesting it reflected NEP1R1 mislocalization rather than altered ER/NE morphology. CTDNEP1^mGFP^ also primarily colocalized with Calnexin in control fat body cells (Fig. 6C). In contrast, CTDNEP1^mGFP^ strongly concentrated at the NE in the *dTorsin^KO^*. This NE-localized CTDNEP1^mGFP^ signal far exceeded the NE-levels of Calnexin (Fig. 6D, arrows). Moreover, while CTDNEP1^mGFP^ was also at the NE of control cells, it appeared uniform around the nucleus (Fig. 6E). CTDNEP1^mGFP^ signal in the *dTorsin^KO^* NE was instead punctate (Fig. 6F). To confirm that Torsin controls CTDNEP1 localization, we examined CTDNEP1^scarlet^ transiently expressed in control mouse embryonic fibroblasts (MEFs), or MEFs that lack TorsinA and TorsinB (Fig. S3G). CTDNEP1^scarlet^ signal was diffuse and colocalized with Calreticulin in control MEFs (Fig. 6G). In contrast, CTDNEP1^scarlet^ was punctate and NE-localized in TorsinA^KO^/TorsinB^KO^ MEFs (Fig. 6H), fully consistent with findings in fly.

**Figure 6:**
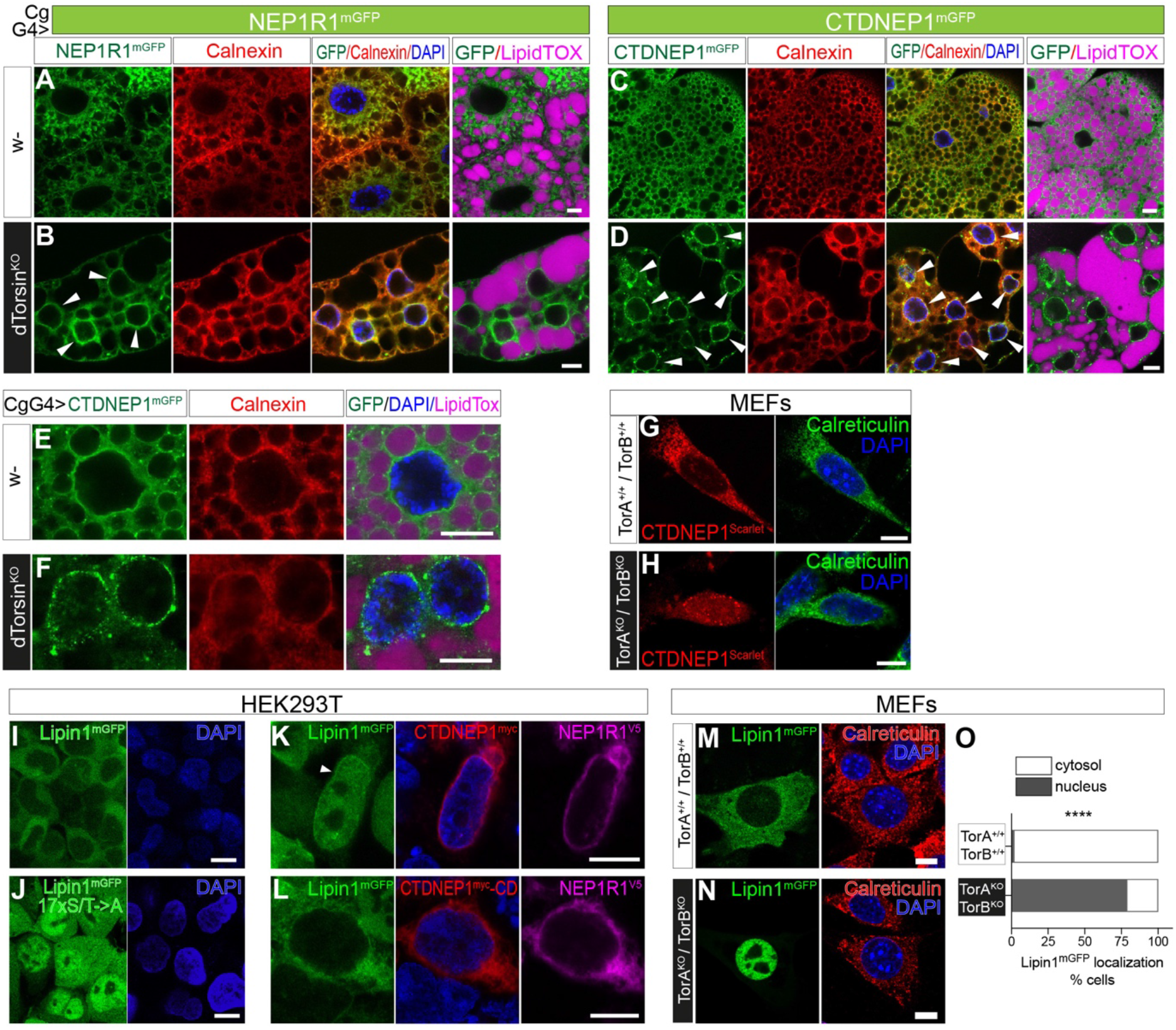
Torsins control the ER/NE localization of CTDNEP1, and the cytosolic/nuclear localization of Lipin. A - F) Control and *dTorsin^KO^* 5DO fat body cells expressing GFP fusion proteins, and labeled with anti-Calnexin (red), and a neutral lipid dye (LipidTOX, magenta). Note different magnifications. All scale bars, 10μm. G & H) Human CTDNEP1^Scarlet^ in MEFs colabeled with the Calreticulin ER marker. I & J) Mouse Lipin1^GFP^-WT and Lipin1^GFP^-17xS/T->A stably expressed in HEK293T cells. K & L) HEK293T-Lipin1^GFP^-WT cells transfected with human CTDNEP1^m^y^c^ / CTDNEP1^myc^-CD (red) and human NEP1R1^V5^ (magenta). Arrow highlights NE enrichment in a transfected cell. M & N) Mouse Lipin1^GFP^ in MEFs colabeled with Calreticulin O) Percentage of MEFs with primarily cytosolic or nuclear Lipin1^mGFP^ signal. **** indicates significant difference (Fisher’s exact test).

Fly Lipin^V5-His^ is exclusively in the cytosol of wild-type fat body cells, but also in the nucleus of *dTorsin^KO^* cells ((Grillet et al., 2016); Fig. S3D & E)). Few studies have examined fly Lipin, but mammalian Lipin1 is known to relocalise to the nucleus when dephosphorylated: a potential explanation for why Torsin loss affects NPC insertion (Han et al., 2012; Peterson et al., 2011). We confirmed wild-type mouse Lipin1^mGFP^ was in the cytosol, while a Lipin1^mGFP^ mutant with 17 serine and threonine residues replaced with alanine (thus suppressing phosphorylation) was in the nucleus (Fig. 6I & J; (Peterson et al., 2011)). NEP1R1^V5^-CTDNEP1^myc^ co-overexpression was sufficient to relocalize Lipin1^mGFP^-WT to the nucleus (Fig. S3F). Indeed, we detected Lipin1^mGFP^ at the nuclear periphery consistent with Lipin1 actively metabolizing PA on the INM (Fig. 6K; arrow). This relocalization did not occur when NEP1R1^V5^ was co-expressed with catalytically dead CTDNEP1^myc^-CD (Fig. 6L; Fig. S3F; (Han et al., 2012)). Finally, we tested whether Torsins control Lipin localization, consistent with them normally suppressing NEP1R1-CTDNEP1 function. Indeed, while Lipin1^mGFP^ was cytosolic in wild-type MEFs, it was exclusively in the nucleus of TorsinA^KO^/TorsinB^KO^ MEFs (Fig. 6M - O). We therefore conclude that Torsins prevent CTDNEP1 from localizing in the NE, and thus prevent Lipin from metabolizing PA at the INM. This potentially explains the *Nep1r1-Ctdnep1* gain-of-function that inhibits NPC insertion in the *dTorsin^KO^*.

### Excess NEP1R1-CTDNEP1/ Lipin activity impairs NPC maturation

We took an overexpression approach to further examine the impact of excess *Nep1r1/ Ctdnep1/ Lipin* activity on the NE. We overexpressed CTDNEP1^mGFP^ or CTDNEP1^mGFP^-CD using the high level r4-Gal4 fat body driver. Overexpressed CTDNEP1^mGFP^ and CTDNEP1-CD^mGFP^ were indistinguishable. They primarily localized in the ER, with some NE localization although this was less than in the *dTorsin^KO^* (Fig. S4A). Megator/Tpr labeled the NE in all cells (Fig. 7A-C). In contrast, the intense mAb414 NE labeling of control cells (Fig. 7D) was absent from cells overexpressing CTDNEP1^mGFP^ (Fig. 7E), but not those overexpressing catalytically dead CTDNEP1^mGFP^-CD (Fig. 7F). Quantification of mAb414 fluorescence confirmed this, as well as showing there was less NE-localized mAb414 labeling in cells overexpressing NEP1R1^mGFP^ and Lipin^V5-His^ compared to ^mGFP^Sec61b (control) and CTDNEP1^mGFP^-CD (Fig. S4B & C).

**Figure 7.**
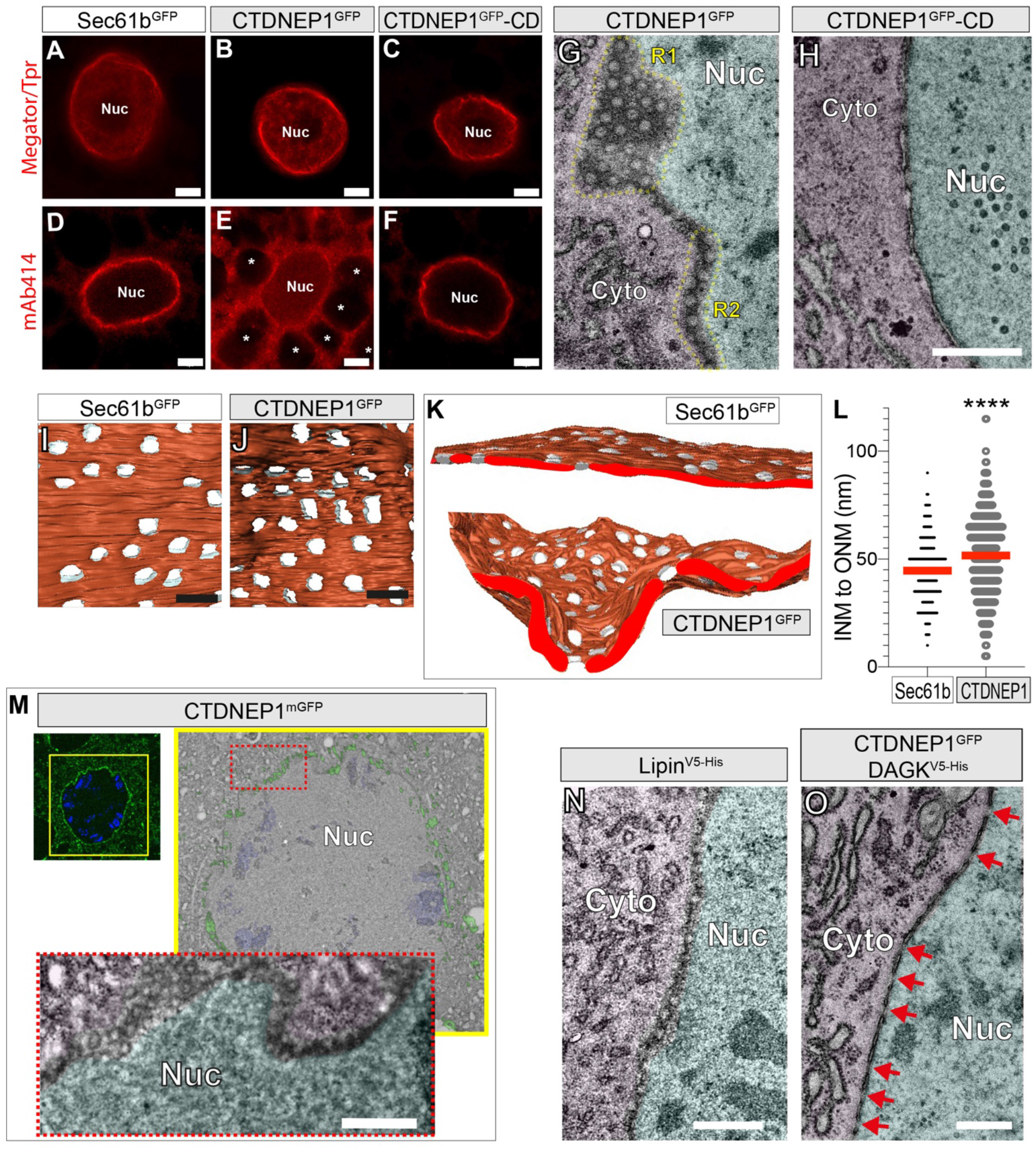
CTDNEP1 phosphatase activity inhibits NPC maturation by dysregulating lipid metabolism. A - F) Fat body nuclei expressing UAS-transgenes with r4-Gal4, and labeled with anti-Megator/Tpr or mAb414. * LD. Scale bars, 5μm. G & H) NE ultrastructure of 5DO fat body cells expressing UAS-transgenes with r4-Gal4. Scale bar, 500nm. I - K) 3D reconstructions of the NE from FIB-SEM images made through a control fat body nucleus (expressing Sec61b) or a NE with CTDNEP1^mGFP^ signal (fluorescence not shown). L) Distance between the INM and ONM at sites immediately adjacent to pores/channels. M) CLEM shows relationship between NE localized CTDNEP1^mGFP^ and NE structure. Left, Super-resolution detection of CTDNEP1^mGFP^; yellow box shows the region examined by SEM. Red box shows enlarged region that contains NE-localized CTDNEP1^mGFP^. Scale bar, 500nm. N & O) NE ultrastructure in 5DO fat body cells expressing UAS-transgenes with r4-Gal4. Scale bar, 500nm.

TEM of CTDNEP1^mGFP^ overexpressing cells detected regions containing circular membrane profiles between a separated INM and ONM (Fig. 7G, R1). Other NE regions (bisected with a different angle of sectioning) had abnormal hemisphere-like profiles that were continuous with the cytosol or nucleus (Fig. 7G, R2). These were indistinguishable to the defects we had detected in *Torip*^-/-^ cells (Fig. 4P). However, they were considerably more frequent upon CTDNEP1^mGFP^ overexpression, and they covered the majority of the NE. No abnormal profiles were seen in CTDNEP1^mGFP^-CD overexpressing cells (Fig. 7H). We used 3D FIB-SEM to define NE structure in CTDNEP1 overexpressing cells. Their NE had regions with abnormally high concave and convex curvature. 3D reconstruction determined that the circular membrane profiles were channels. They were present at a similar density to nuclear pores in control cells (Fig. 7I vs J), but the INM and ONM adjacent to channels were significantly further apart than they were at the NPC of control cells (Fig. 7K & L).

We tested whether excess NE-localized PA metabolism caused these NPC defects. Using Correlative Light and Electron Microscopy (CLEM), we identified that they appeared at sites where CTDNEP1^mGFP^ accumulated in the NE (Fig. 7M). Lipin^V5-His^ overexpression also produced these abnormal structures (Fig. 7N). Next, we used DAG kinase (DAGK) as a tool to establish NPC are affected by altered lipid levels, and not the accumulation of CTDNEP1 or Lipin proteins. DAGK opposes Lipin by converting DAG to PA, and localizes in the nucleus as well as the cytosol (Fig. S4D). NE structure was normal in DAGK^V5-His^ expressing cells (Fig. S4E vs F). It was also normal in cells co-expressing DAGK^V5-His^ and CTDNEP1^mGFP^ (Fig. 7O), while cells co-expressing Sec61b with CTDNEP1^mGFP^ continued to develop NE abnormalities (Fig. S4G vs H). Taken together we conclude that NEP1R1-CTDNEP1/ Lipin control of PA and DAG has a previously unrecognized importance for NPC maturation.

## Discussion

These data show that Torsin, NEP1R1-CTDNEP1 and Lipin form a molecular pathway that couples the ER/ NE lumen to PA and DAG levels on the opposite face of the membrane. Torsins suppress pathway activity, while NEP1R1-CTDNEP1 promotes Lipin-mediated PA conversion to DAG. We show that this pathway has sufficient control over PA metabolism that it broadly affects the cell, including controlling growth, organelle size, LD biogenesis and nuclear membrane structure. Pathway regulation is especially active at the NE, where it is required for two distinct steps of NPC biogenesis that were not previously known to be influenced by lipids.

There is accumulating evidence that the INM is a major site of cellular lipid metabolism. Lipin has an NLS, NEP1R1 and CTDNEP1 are named after their NE localization, and the CCT rate limiting enzyme of membrane lipid synthesis also localizes in the nucleus (Wang et al., 1993). It has been shown that PA conversion to DAG and onto TAG takes place on the yeast INM (Romanauska and Kohler, 2018). Our data now show this is also the case in animal cells given the nuclear TAG that appears with combined *dTorsin* and *Torip* loss. The reasons for INM-localized lipid metabolism are only just becoming clear. It is important for mitotic nuclear growth and nuclear reformation (Makarova et al., 2016; Penfield et al., 2020). It is potentially also advantageous to segregate lipid reactions away from the bulk ER, and thus prevent newly synthesized lipids from inducing stress and protein misfolding (Jacquemyn et al., 2017). However, we now identify that NE-localized lipid metabolism can have negative consequences, and is under the control of Torsins and the structurally related Torip/LAP1.

The first negative consequence we identified is that deregulated PA metabolism interferes with the INM/ONM fusion of NPC insertion. We make this conclusion since 1) Torsin concomitantly excludes Lipin from the nucleus and is required for NPC biogenesis, and 2) that NEP1R1-CTDNEP1 overactivity underlies why NPC insertion fails in the *dTorsin^KO^*. Additionally, we previously linked excess Lipin PA metabolism to Torsin-membrane defects in interphase mammalian neurons (Cascalho et al., 2020), and note that similar INM herniations are associated with excess PA-DAG-TAG metabolism at the INM of yeast (Romanauska and Kohler, 2018). Additional work is needed to uncover why excess PA to DAG metabolism prevents INM/ONM fusion. Several scenarios are possible. One is that PA or DAG recruit fusion proteins to the emerging NPC, and this fails with too little or too broadly distributed lipid. Indeed, PA has been shown to recruit ESCRTs to poorly inserted NPC (Thaller et al., 2020) – although we note that the almost complete absence of normal NPC in the *dTorsin^KO^* can only be explained by impaired biogenesis. Another possibility is that PA metabolism affects biophysical membrane properties beyond what is permissive for fusion. This is consistent with the fact that PA metabolism affects many lipid classes (Grillet et al., 2016), and that NPC insertion in yeast is sensitive to membrane fluidity/ deformability (Hodge et al., 2010; Lone et al., 2015).

We additionally also identify that excess INM-localized PA to DAG conversion prevents a later step of NPC biogenesis. We uncovered this in cells overexpressing NEP1R1, CTDNEP1, or Lipin, and the same NPC defects also appeared in *Torip*^-/-^ cells. All read-outs indicate that these perturbations more mildly affect NE lipid metabolism compared with the hyperstimulation of Torsin loss. This can explain why INM/ONM fusion occurs. Nevertheless, abnormal NE PA metabolism prevents NPC maturation, leading to nucleo-cytoplasmic channels that are substantially longer than a mature NPC, and lack channel FG nucleoporins. This type of defect has not been previously described; thus, again, additional work is required to define what aspect of NPC maturation is blocked. One possibility is that structural nucleoporins are recruited by PA or DAG and, in turn, their absence prevents INM/ONM compression and channel nucleoporin recruitment. The larval fat body will be ideal to further explore this question, as well as to dissect other events of interphase NPC biogenesis that remains poorly defined compared to mitotic NPC insertion.

Torsins are AAA+ enzymes inside the ER/NE lumen. AAA+ enzymes typically operate as oligomeric rings that use the energy of ATP hydrolysis to alter the structure of binding partners. This can include disassembly of otherwise stable protein complexes (Hanson and Whiteheart, 2005). CTDNEP1 is the transmembrane phosphatase, while NEP1R1 is the regulatory unit that stabilizes CTDNEP1 against degradation (Han et al., 2012). Heterodimeric NEP1R1-CTDNEP1 has a conserved transmembrane topology, meaning it is accessible to Torsins. Thus, the simplest explanation for our findings is that Torsins engage the luminal domains of NEP1R1-CTDNEP1, resulting in complex dissociation, less CTDNEP1 phosphatase activity and, in turn, less Lipin mediated PA to DAG conversion. The next steps are to determine whether Torsin homomers indeed directly drive NEP1R1-CTDNEP1 complex disassembly. In addition, our data strongly argue that Torsins do not act as classic AAA+ enzymes. Instead, Torsin homo-oligomerization into ATPase inactive structures is key to NEP1R1-CTDNEP1/Lipin pathway suppression. This includes that Torsin continues to suppress the pathway when Torip is absent, and we genetically prove that *Torip* and *Torsin* independently and similarly affect lipid metabolism.

Finally, Torsins and Torip/LAP1 have clinical relevance because complete *TOR1A/TorsinA* loss causes a congenital neurological syndrome (Kariminejad et al., 2017), complete Torip/LAP1 loss causes multisystemic congenital disease (Fichtman et al., 2019), and a heterozygous *TOR1A*/TorsinA mutation causes the neurological movement disorder of dystonia (Ozelius et al., 1997). We have previously shown that Lipin inhibition is beneficial in *TOR1A* disease mouse models (Cascalho et al., 2020). Here we now show that *Nep1r1* and *Ctdnep1* inhibition completely prevent the broad cellular defects of Torsin loss. This includes defects with clear relevance for *TOR1A*/TorsinA and LAP1 diseases: the reduced ER capacity, excess PA metabolism, excess TAG/LD deposition, abnormal NPC, and defective NE structures that we normalize in fly, have all been described in mammalian *TOR1A/ TOR1AIP1* models or patient cells (Cascalho et al., 2020; Fichtman et al., 2019; Goodchild et al., 2005; Hewett et al., 2007; Kim et al., 2010; Pappas et al., 2018; Shin et al., 2019). We therefore conclude that NEP1R1-CTDNEP1/Lipin pathway overactivity 1) indeed underlies the broad and diverse defects of Torsin loss, 2) that abnormal lipid metabolism likely underlies this series of genetic human neurological and neuromuscular diseases, and 3) that NEP1R1 and CTDNEP1, as well as Lipin, should be advanced as therapeutic targets.

## Acknowledgments

This work was only possible by the broad support provided by the Foundation for Dystonia Research. J.J, and J.F, have FWO Sb fellowships (1S07317N and 1S54121N). We thank the Dystonia Medical Research Foundation for an award. Data storage was supported by an ISPAMM project grant (AKUL/13/39). The Nikon A1R Eclipse Ti was acquired through a Hercules type I grant (AKUL/09/037) to Wim Annaert. We make a major acknowledgement to Jens Rummens for insightful discussions and making cell lines, to Nils Schoovaerts for plasmids, and to visiting students Noémie De Broeck, Sara Garcia Garotte and Laura Ordonez Cuerva. We thank the Gent VIB EM facility, especially Anneke Kremer, and the Leuven VIB Light Microscopy Expertise Unit, especially Benjamin Pavie. Ragna Sannerud and the Annaert Lab, as well as the Schmucker Lab, gave advice on generating cell lines.

## Author contributions

J.J. and R.E.G. were involved in project conceptualization, data collection and analysis, supervised experiments, prepared figures and prepared the manuscript. R.E.G. and J.J. equally share copyright. J.F., J.J., S.F.G., J.S., K.V. performed experiments. P.V. and N.V.G. supervised EM experiments.

