## Supplementary material for "The Torsin/ NEP1R1-CTDNEP1/ Lipin axis regulates nuclear envelope lipid metabolism for nuclear pore complex insertion": Table S3

**Supplemental Table S3**

| Antibody | Concentration | Vendor/ Reference |
|---|---|---|
| mAb414 | 1:50 | Abcam #24609 |
| Anti Megator/Tpr (*Drosophila)* | 1:100 | DSHB 12F10-5F11 |
| Anti Lam*Dm*0 | 1:30 | DSHB ADL101 |
| Anti V5 | 1:100 (fly) and 1:1000 (cells) | Life Technologies R960-25 |
| Anti c-Myc | 1:125 | Invitrogen #13-2500 |
| Anti Calnexin (*Drosophila)* | 1:30 | DSHB Cnx99A 6-2-1 |
| Anti TorsinB (mammalian) | 1:500 | {Jungwirth, 2010 #458} |
| Anti $\alpha$-Tubulin (mammalian) | 1:10.000 | Sigma |
| Anti Calreticulin | 1:500 | Sigma #C4606 |
| Anti Mouse - Alexa Fluor 488 | 1:100 | Jackson #715-545-151 |
| Anti Mouse - rhodamine red | 1:100 | Jackson #715-295-150 |
| Anti Mouse - Cy5 | 1:100 | Jackson #715-175-151 |
| Anti Rabbit HRP | 1:5000 | Jackson |
| Anti Mouse HRP | 1:5000 | Jackson |
