## Supplementary material for "The Torsin/ NEP1R1-CTDNEP1/ Lipin axis regulates nuclear envelope lipid metabolism for nuclear pore complex insertion": Table S1

**Table1: Drosophila stocks and transgenic fly lines**

| Line name | Genotype | Stock number (if relevant) | Reference (if relevant) |
|---|---|---|---|
| Control | w1118 | BDSC (3605) | |
| Genomic insertions and deletions | | | |
| *dTorsin$^{KO}$* | w-, dTorsinKO78/ FM7i, Act-GFP | | (Wakabayashi-Ito et al., 2011) |
| *Torip (CG14103)* | w1118;;Torip(CG14103) 11ntΔ/TM6C,tb | | This study |
| *GFP-Nup358* | emGFP::nup358 PBac{y[+mDint2] = vas-Cas9}VK00027 | | (Hampoelz et al., 2019) |
| GAL4 lines | | | |
| CgG4 | w1118; Cg-GAL4 | BDSC (7011) | (Hennig et al., 2006) |
| r4G4 | W1118;; r4-GAL4 | BDSC (33832) | (Lee and Park, 2004) |
| RNAi lines | | | |
| *LipinLOF* | yw; Lipin KG00562/CyO | BDSC (13293) | (Pereira et al., 2011) |
| *Wunen1* | w1118;; UAS-Wunen1 RNAi/TM3 | VDRC (v6446), rebalanced over TM6C,tb in our lab | (Zhang et al., 1996) |

| | | | |
|---|---|---|---|
| *Wunen2* | w1118;; UAS-Wunen2 RNAi | VDRC (v4176), rebalanced over TM6C,tb in our lab | (Kraut et al., 2001) |
| *Agpat2 (CG17608)* | w1118; UAS-AgpatRNAi/CyO | VDRC (v51162) | (Kong et al., 2010) |
| *Gpat4* | w1118;; UAS-GpatRNAi/TM3 | VDRC (v10281), rebalanced over TM6C,tb in our lab | (Wilfling et al., 2013) |
| *Fitm* | w1118; UAS-FitmRNAi/CyO | BDSC (57431) | (Kadereit et al., 2008) |
| *Seipin* | w1118; UAS-SeipinRNAi/CyO | BDSC (37501) | (Tian et al., 2011) |
| *Ddk* | w1118; UAS-DGKRNAi/CyO | VDRC (v38239), rebalanced over Cyo,tb in our lab | (Trinh et al., 2019) |
| *Ctdnepl* | w1118; UAS-DdRNAi/CyO | VDRC (v12941) | (Liu et al., 2011) |
| *Neplrl (CG41106)* | w1118; UAS-NeplrlRNAi/CyO | VDRC (v103171) | |
| *Neplr2 (CG8009)* | w1118; UAS-Neplr2RNAi/CyO | VDRC (v48955) | (Schnorrer et al., 2010) |
| *Koi* | w1118; UAS-KoiRNAi/CyO | BDSC (40924) | (Kracklauer et al., 2007) |
| *Lipin* | w1118; UAS-LipinRNAi | VDRC (v36006) | (Ugrankar et al., 2011) |
| *Luciferase* | w1118;; UAS-LuciferaseRNAi | BDSC (31603) | (Perkins et al., 2010) |

| UAS lines | | | |
|---|---|---|---|
| UAS-dTorsin$^{mGFP}$ | w1118; UAS-dTorsin-mGFP/CyO | | (Grillet et al., 2016) |
| UAS-Torip$^{mGFP}$ | w1118; UAS-CG14103-mGFP/CyO | | (Grillet et al., 2016) |

| | | | |
|---|---|---|---|
| UAS-$^{dTomato}$Sec61b | w1118; UAS-dTomato-Sec61β/CyO | BDSC (64746) | (Summerville et al., 2016) |
| UAS-dTorsin$^{mGFP}$-K114A- | w1118; UAS-dTorsin-K114A-mGFP /CyO | Attp40 | In this study |
| UAS-dTorsin$^{mGFP}$-E177Q | w1118; UAS-dTorsin-E177Q-mGFP/CyO | Attp40 | In this study |
| UAS-dTorsin$^{mGFP}$-G256D | w1118; UAS-dTorsin-G256D-mGFP /CyO | Attp40 | In this study |
| UAS-$^{mGFP}$Sec61b | w1118; UAS-Sec61β-mGFP /CyO | Attp40 | In this study |
| UAS-CtdnepI$^{mGFP}$ | w1118; UAS- CtdnepI-mGFP /CyO | Attp40 | In this study |
| UAS-NepIrl$^{mGFP}$ | w1118; UAS- NepIrl-mGFP /CyO | Attp40 | In this study |
| UAS-CtdnepI$^{mGFP}$-CD | w1118; UAS- CtdnepI-CD-mGFP /CyO | Attp40 | In this study |
| UAS-Lipin$^{V5\text{-}His}$ | w1118;; UAS- Lipin$^{V5\text{-}His}$ /TM6b, tb | Attp40 | In this study |
| UAS-Dgk$^{V5\text{-}His}$ | w1118; UAS- Dgk$^{V5\text{-}His}$ /CyO or TM6b,tb | Attp40 | In this study |
