## Supplementary material for "The Torsin/ NEP1R1-CTDNEP1/ Lipin axis regulates nuclear envelope lipid metabolism for nuclear pore complex insertion": Table S2

**Table2: Primers used in this study**

| Gene | qRT-PCR |
| --- | --- |
| *Torip* R1 | 5’TCACAACGGCATCGGTATTA3’/ 5’CCTGGTGCAGGAACAAAAAT3’ |
| *Torip* R2 | 5’CGAGGGGCTGATTAACAAAA3’/ 5’TGAATTGTTCAATCGCCAAA3’ |
| *Torip* R3 | 5’ATACGGTTTGGCGATTGAAC3’/ 5’CAACTCCTGGTCCCACAGAT3’ |
| *Ctdnep1* | 5’GCCAAGTGCGAGCTTTTATC3’/ 5’GATTCCGATCGATGGTCACT3’ |
| *Nep1r1* | 5’TGGTTTTCCAAATCCTGTCC3’/ 5’ATCAATGTTAAGGCGGAACG3’ |
| *Lipin* | 5’GTCGCAACCTAAGCGAAAAG3’/ 5’TTGTGACCGTGGTGAGGTTA3’ |
| *RpL32* | 5’TACAGGCCCAAGATCGTGAA3’/ 5’GTTCGATCCGTAACCGATGT3’ |
| *Tor1b* | 5’CTTGCCACCGAAGTGATT3’/ 5’CAAGGACAGAGTCAATGGTTT3’ |
| *Pgk1* | 5’TTC CCA TGC CTG ACA AGT3’/ 5’CCCAGCAGAGATTTGAGTTC3’ |
| *Hprt1* | 5’AAT CAT TAT GCC GAG GAT TTG G3’/ 5’GAGCAAGTCTTTCAGTCCTGT3’ |
| *B2m* | 5’TCT GGT GCT TGT CTC ACT3’/5’TTCGGCTTCCCATTCTCC3’ |

| Gene | Genomic Locus Amplification |
| --- | --- |
| *Torip* | 5’ AAAGCTACGAGGAAGGGA 3’/ 5’ CAAATAGAACACCGGCAA 3’ |
| *Ctdnep1-mGFP* | 5’ CAACGCCATACCCATCAA 3’/ 5’CCTCAGTGCTCCTTCTAA 3’ |
| *Tor1b* | 5’ GTCTCCTGTGTGTGTGTG 3’/ 5’ TTCAGTTCTCCAGGTGGT 3’ |

| UAS Construct | Fragment Amplification |
|---|---|
| pUAST$^{V5\text{-}His}$ | 5′ GCCGCGGCTCGAGGGTACCTGGCCGGCTGGGCCCGTTTCG 3′<br>5′ GGTTCCTTCACAAAGATCCTTCAATGGTGATGGTGATGATGACCGGTACGCG 3′ |
| dTorsin$^{mGFP}$-G256D | <u>5’ fragment (N-terminal dTorsin)</u><br>5'ACTCTGAATAGGGAATTGGGATGCACGCATGTTATCACTGTGCC 3'<br>5' CTTTTTCATATCGCCATCCATGTTATAGGCGGCCTTCC 3′<br><u>3’ fragment (C-terminus and mGFP)</u><br>5' ATGGATGGCGATATGAAAAAGACCACCATGATTGAGTCTCATGTTATCG 3'<br>5' AAGATCCTCTAGAGGTACCCTTACTTGTACAGCTCGTCCATGCCGAGAG 3' |
| dTorsin$^{mGFP}$-E177Q | <u>5’ fragment (N-terminal dTorsin)</u><br>5' ACTCTGAATAGGGAATTGGGATGATGAGCTTTCCACGCATGTTATCACTGTGCC 3'<br>5' TTATCCACCTGGTCGAAGATGAACAAAGATCTCGGG 3'<br><u>3’ fragment (C-terminus and mGFP)</u><br>5' CATCTTCGACCAGGTGGATAAAATGCCCAGC 3'<br>5' AAGATCCTCTAGAGGTACCCTTACTTGTACAGCTCGTCCATGCCGAGAGTG 3' |
| dTorsin$^{mGFP}$-K114A | <u>5’ fragment (N-terminal dTorsin)</u><br>5' ACTCTGAATAGGGAATTGGGATGATGAGCTTTCCACGCATGTTATCACTGTGCC 3'<br>5' CACAAAATTCGCGCCCGTACCGGGCTGG 3'<br><u>3’ fragment (C-terminus and mGFP)</u><br>5' CGGTACGGGCGCGAATTTTGTGGCTGAGCAAATTGC 3'<br>5' AAGATCCTCTAGAGGTACCCTTACTTGTACAGCTCGTCCATGCCGAGAGTG 3' |

| | |
|---|---|
| CTDNEP1$^{mGFP}$ | 5’ fragment (CTDNEP1)<br>5’ CTCTGAATAGGGAATTGGGCAAAATGATTTCGCTGCTGCAAATGAA 3’<br>5’ CTCACCATCCAGAGGCGGTGCAGG 3’<br>3’ fragment (mGFP)<br>5’ GCCTCTGGATGGTGAGCAAGGGCGAG 3’<br>5’ GATCCTCTAGAGGTACCCTTACTTGTACAGCTCGTCCATGCC 3’ |
| CTDNEP1$^{mGFP}$-CD | 5’ fragment (N-terminal CTDNEP1)<br>5’ ACTCTGAATAGGGAATTGGGATGATTTCGCTGCTGCAAATGAAATTCCGTGC 3’<br>5’ TAGCGTTTCCTCCAGCTCCAGAACGAGGGTCTTGCGCTGC 3’<br>3’ fragment (C-terminal CTDNEP1 and mGFP)<br>5’ TCGTTCTGGAGCTGGAGGAAACGCTAATCCACTCCCATCACAATGCG 3’<br>5’ AAGATCCTCTAGAGGTACCCTTACTTGTACAGCTCGTCCATGCCGAGAG 3’ |
| NEP1R1$^{mGFP}$ | 5’ fragment (NEP1R1)<br>5' CTCTGAATAGGGAATTGGGCAAAATGGAGGACACAGATCAGGAAG 3'<br>5’ GCTCACCATCTCTGTCTGCCGTGGAAGTAGG 3’<br>3’ fragment (mGFP)<br>5' AGACAGAGATGGTGAGCAAGGGCGAG 3'<br>5' GATCCTCTAGAGGTACCCTTACTTGTACAGCTCGTCCATGCC 3' |
| DAGK$^{V5\text{-}His}$ | 5’ ACTCTGAATAGGGAATTGGGGCGTCCCGATCCGCACACATGAGATGG 3’<br>5’ GCCCAGCCGGCCAGGTACCCGCTGCACAGCAGGTTGAAAAATCCGCGTC 3’ |
| Lipin$^{V5\text{-}His}$ (isoform A) | 5’ ACTCTGAATAGGGAATTGGGATGAATAGCCTGGCGCGGGTTTTCAG 3’<br>5’ GCCCAGCCGGCCAGGTACCCGGAATTAATCCACTTGGGAGGCGAAGCC 3’ |

| | |
|---|---|
| $^{mGFP}$Sec61b | <u>5’ fragment (mGFP)</u><br>5’ ACTCTGAATAGGGAATTGGGATGGTGAGCAAGGGCGAGGAGCTG 3’<br>5’ TGGCATCAGCCTTGTACAGCTCGTCCATGCCGAGAGTG 3’<br><u>3’ fragment (Sec61b)</u><br>5’ GCTGTACAAGGCTGATGCCAAGCTAGCAATGCCTGG 3’<br>5’ AAGATCCTCTAGAGGTACCCTTACGAACGAGTGTACTTGCCCCAAATGTGC 3’ |

| Mammalian Construct | Fragment Amplification |
|---|---|
| pcDNA5⁄FRT⁄TO-Lipin1$^{mGFP}$<br>and<br>pcDNA5⁄FRT⁄TO-Lipin1$^{mGFP}$-17xS/T->A | <u>5’ fragment (mouse Lipin1)</u><br>5’ GACTCTAGCGTTTAAACTTAATGGAAATGGACTACAAGGATGACGATGACAAGGG 3’<br>5’ GCGGTACCGTAGCTGAGGCTGAATGCATGTCCTGG 3’<br><u>3’ fragment (linker + mGFP)</u><br>5’ AGCCTCAGCTACGGTACCGCGGGCCCGG 3’<br>5’ CTGGAACATCATATGGATACTTACTTGTACAGCTCGTCCATGCCGAGAGTGATCC 3’ |
| pBudCE4.1-CTDNEP1$^{myc\text{-}His}$/ NEP1R1$^{V5\text{-}His}$ | <u>CTDNEP1 (no stop codon)</u><br>5' ACCCAAGCTTATGATGCGGACGCAGTGTCTG 3'<br>5' ATCCTCTAGAAGTACTGCTTGACCAGAGCCGATGTTGGTGAAG 3’<br><u>NEP1R1 (no stop codon)</u><br>5' CGTGAACACGTGGTCGCGGCATGAACTCGCTGGAGCAGG 3'<br>5' GATCTCTCGATTGAACATGAGGCCTAGGTTTC 3'<br><u>Plasmid backbone (pCMV)</u><br>5' TCATGTTCAATCGAGAGATCTGGCCGGC 3'<br>5' TCCGCATCATAAGCTTGGGTCTCCCTATAGTGAGTCG 3' |

| | |
|---|---|
| | Plasmid backbone (pEF1a)<br>5' AAGCAGTACTTCTAGAGGATCCGAAC 3'<br>5' GCCGCGACCACGTGTTC 3' |
| pBudCE4.1-CTDNEP1$^{\text{myc-His}}$-CD/ NEP1R1$^{\text{V5-His}}$ | 5' CTDNEP1 fragment (N-terminal)<br>5' ACTCACTATAGGGAGACCCAATGATGCGGACGCAGTGTCTGCTGG 3'<br>5' TAAGTGTCTCTTCCAGCTCCAGCACCAGGATCTTCCTCTTCACC 3'<br>3' CTDNEP1 fragment (C-terminal)<br>5' TGGTGCTGGAGCTGGAAGAGACACTTATTCACTCCCACCATGATGGGG 3'<br>5' TGAGTTTTTGTTCGGATCCTGAAGTACTGCTTGACCAGAGCCGATGTTGG 3' |
| pBudCE4.1-CTDNEP1$^{\text{Scarlet}}$ | Scarlet (step 1)<br>5' GTCAAGCAGTACTTCTAGAGTCGCCACCATGGTGAGCAAGGGC 3'<br>5' CTGAGATGAGTTTTTGTTCGCTACTTGTACAGCTCGTCCATGCCGCCG 3'<br>CTDNEP1$^{\text{Scarlet}}$ (step 2) into pBudCE4.1<br>5' AAGGTACCAGCACAGTGGACATGATGCGGACGCAGTGTCTGCTGG 3'<br>5' GAAACGGGCCCAGCCGGCCACTACTTGTACAGCTCGTCCATGCCGCC 3' |
| pBudCE4.1-NEP1R1$^{\text{Scarlet}}$ | Scarlet (step 1)<br>5' ATCGAGAGATCTGGCCGGCTTCGCCACCATGGTGAGCAAGGGC 3'<br>5' ACCTTCGAAACGGGCCCAGCCTACTTGTACAGCTCGTCCATGCCGCCG 3'<br>NEP1R1$^{\text{Scarlet}}$ (step 2) into pBudCE4.1<br>5' AAGGTACCAGCACAGTGGACATGAACTCGCTGGAGCAGGCGGAAG 3'<br>5' GAAACGGGCCCAGCCGGCCACTACTTGTACAGCTCGTCCATGCCGCC 3' |
