## Supplementary material for "The Torsin/ NEP1R1-CTDNEP1/ Lipin axis regulates nuclear envelope lipid metabolism for nuclear pore complex insertion": Materials and Methods

### Fly Stocks and maintenance

All *Drosophila melanogaster* stocks (Table S1) were handled using standard protocols, fed on a standard diet consisting of cornmeal, agar, yeast, sucrose, and dextrose and kept under a 12 hr light/dark cycle. All experimental crosses were kept at 25°C.

### Molecular Biology

#### *New Drosophila Lines*

Microinjections were performed by BestGene Inc. (CA, USA). We used CRISPR to disrupt genomic sequences by injecting vas-cas9 (X) flies (Norrander et al., 1983) with pCFD3-dU6:3gRNA (Addgene #49410; (Port et al., 2014)) carrying guide RNA sequences. Transgenic UAS fly lines were produced by injecting pUAST plasmids into Attp2+ or Attp40+ stocks for PhiC31 integrase-mediated site specific transgenesis.

#### *Torip$^{KO}$ line*

The flyCRISPR Target finder Tool (https://flycrispr.org/) predicted a guide RNA 5'-gCCGATCAGTCCAGAAGACG-3' that would introduce a double stranded break into *Torip* exon one, 191bp after the start codon. $G_0$ flies were crossed with w/y; *Tub*-GAL4/TM6C, Tb flies. 158 offspring with white eyes and short bristles were expanded into lines. The *Torip* locus of each line was PCR amplified (Table S2) and sequenced to identify lines where non-homologous end joining introduced frameshift mutations. We identified two lines with an 11-nucleotide deletion (Δ11). Homozygous *Torip* Δ11 males and females were viable and fertile.

*UAS constructs*

One or more cDNA sequences were cloned between the EcoRI and XhoI sites of pUAST-Attb (Bischof et al., 2007) using the Gibson NEBuilder® Hifi DNA assembly kit. Primer sequences, including overhangs for Gibson cloning, are provided in Table S2. We also generated a modified pUAST_Attb-V5-His backbone by assembling a 135bp PCR amplified fragment containing the V5 and His tag from pBudCE4.1 (Invitrogen™, V53220) with pUASTAttb linearized with XbaI. The coding region of all pUAST plasmids were fully verified by Sanger sequencing before further use.

pUAST-mGFP-Sec61b was produced by assembling pUAST with 1) a 5’ PCR fragment encoding mGFP (amplified from pCR8-dTorsin-mGFP) and 2) a 3’ PCR fragment encoding mouse Sec61b amplified from pcDNA3 SEC61B-V5-APEX (Addgene #83411; (Lee et al., 2016)).

pUAST-dTorsin plasmids with mutagenized dTorsin were produced by assembling two PCR-amplified DNA fragments into pUAST. Both reactions used pUAST-dTorsin$^{mGFP}$ as the template (Grillet et al., 2016). PCR 1 generated the 5’ sequence from the ATG of dTorsin until the mutation, and PCR 2 generated a 3’ fragment extending from immediately after the mutation to the stop codon of mGFP. 3’ and 5’ primers of PCR 1 and PCR 2, respectively, carried the mutation.

*Ctdnepl (Dd,* NM_134605) and *Neplrl* (*CG41106*, NM_001042844) cDNAs lacking the stop codon were PCR amplified from a third instar larvae cDNA library. These were assembled with a PCR fragment encoding a linker and mGFP. We generated phosphatase-dead CTDNEP1 by mutating the catalytic LDLD motif to LELE (Kim et al., 2007), via assembling two PCR-amplified DNA fragments into pUAST. Both reactions used pUAST-CTDNEP1$^{mGFP}$ as the template. PCR 1 generated the 5’ sequence from the ATG to the site of the mutation, and PCR 2 generated the 3’ sequence extending from immediately after the mutation to the stop codon of mGFP. 3' and 5' primers of PCR 1 and PCR 2, respectively, carried the mutation.

*Dgk (DgK,* DAGK*, NM_078930.4) and Lipin (Lpin, NM_136515)* cDNA sequences lacking the stop codon were amplified from EST clones (BT003323 and GH19076 Source BioScience) and assembled as individual fragments with pUAST_Attb-V5-His.

*Mammalian Expression Plasmids*

All cloning was done with the Gibson NEBuilder® Hifi DNA assembly kit. Primer sequences, including overhangs for Gibson cloning, are provided in Table S2. Mouse Lipin1 and GFP were amplified and inserted into pcDNA5⁄FRT⁄TO (Invitrogen) linearized by HindIII and XhoI, as a three-fragment Gibson cloning reaction. Lipin1 was PCR amplified from "pRK5 FLAG wildtype, catalytic active lipin1" or "pRK5 FLAG 17xS/T->A, catalytic active lipin1" that were gifts from David Sabatini (Addgene plasmid # 32005, 32007 (Peterson et al., 2011)). mGFP with a short linker was amplified from a modified pEGFP-N1 (Clontech).

We produced dual promoter pBudCE4.1-CTDNEP1$^{myc\text{-}His}$/NEP1R1$^{V5\text{-}His}$ as a four fragment Gibson assembly. Human CTDNEP1 was amplified from cDNA clone MGC:16648 IMAGE:4123279, human NEP1R1 was amplified from cDNA clone MGC:41938 IMAGE:5266084, and pBudCE4.1 (Invitrogen) was the template for PCR amplification of the two backbone fragments. CTDNEP1-CD was produced by mutating the catalytic LDLD motif to LELE (Kim et al., 2007). The pBudCE4.1-CTDNEP1$^{myc\text{-}His}$-CD/ NEP1R1$^{V5\text{-}His}$ plasmid was generated by excising wild-type CTDNEP1 from pBudCE4.1-CTDNEP1$^{myc\text{-}His}$/NEP1R1$^{V5\text{-}His}$ as a HindIII and XbaI fragment. The backbone was then re-assembled with CTDNEP1-CD produced as two PCR fragments. CTDNEP1 PCR 1 generated the 5' DNA fragment from the ATG to the site of the mutation, and PCR 2 generated the 3' CTDNEP1 fragment from immediately after the mutation to the penultimate codon. 3' and 5' primers of PCR 1 and PCR 2, respectively, carried the "LELE" mutation. pBudCE4.1-CTDNEP1$^{Scarlet}$ and pBudCE4.1-NEP1R1$^{Scarlet}$ plasmids were produced in two steps. Scarlet sequence was amplified from pmScarlet_H2A_C1 (a gift from Dorus Gadella (Addgene plasmid #85051 (Bindels et al., 2017)) and inserted into pBudCE4.1-CTDNEP1$^{myc\text{-}His}$/ NEP1R1$^{V5\text{-}His}$ linearized with BamH1 (to produce the CTDNEP1 fusion) or SfiI (to produce the NEP1R1 fusion). Then CTDNEP1$^{Scarlet}$ and NEP1R1$^{Scarlet}$ sequences were amplified and re-cloned by Gibson assembly in front of the pEF1$\alpha$ promoter of empty pBudCE4.1 linearized by Xho1 and BglII.

**Fly Tissue Preparation**

Male and female flies were introduced to fresh vials, followed by 12 hours of egg laying, and then the removal of adults. Larvae that were present 3, 4, or 5 days later are referred to as 3 DO, 4 DO, and 5 DO, respectively. Individual larvae were dissected and all tissue expect adipose and nervous system tissue was removed to expose the fat body under a stereomicroscope. Bright field images were collected with a Zeiss Discovery V12 stereomicroscope after all abdominal tissue except adipose and nervous system tissue was removed. Alternatively, preparations were fixed as described below, or the fat body was removed and snap frozen on dry ice.

**qRT-PCR**

Total RNA was extracted from isolated fat bodies or cell pellets using the RNeasy Mini Kit (Qiagen) according to the manufacturer's protocol, followed by DNAseI treatment. Subsequently, cDNA was generated from 1μg of RNA using the SuperScript II First-Strand Synthesis System for RT-PCR (Life Technologies) using a mix of oligo(dt) primers and random hexamer primers. A mock control without reverse transcriptase was included for all samples. Next, 4.5µL of 25 ng/µL cDNA, mock cDNA condition, or water were added to 1.5µL of 5µM forward and reverse primers, and the LightCycler 480 SYBR Green I Master Mix (Roche). Reactions and analysis were performed with a LightCycler 480 (Roche). Primers are listed in Table S2. All qPCR runs were performed in triplicate. Amplification efficiency of each reaction was normalized to reactions performed with primers against the ribosomal house-keeping gene, *RpL32* for *Drosophila* or three housekeeping genes (*Hprt1*, *Pgk1*, *B2m)* for Mouse Embryonic Fibroblasts (MEFs), and relative fold gene expression quantified by the $2^{-\Delta\Delta Ct}$ method.

**Cell Lines**

*Flp-In HEK293T*

HEK293T lines stably expressing mouse Lipin1$^{mGFP}$ were produced using the Invitrogen Flp-In system according to the manufacturer's instructions. Briefly, the Flp-In Host cell line (Flp-In T-Rex 293) was cultured under standard conditions, including 10% certified tetracycline-free FBS, 15 µg/ml blasticidin, and 100 µg/ml zeocin. Cells were co-transfected with pcDNA5⁄FRT⁄TO-Lipin1$^{mGFP}$ or pcDNA5⁄FRT⁄TO-Lipin1$^{mGFP}$-17xS/T->A and pOG44 using Lipofectamine 2000 (ThermoFisher Scientific) following manufacturer's instructions. Transfected cells were expanded in the original media, also including 100 µg/ml hygromycin to establish the new lines. Further experiments were performed with cells plated onto coverslips precoated with 0.1mg/ml Poly–L–lysine (Sigma). These were transfected with pBudCE4.1 plasmids using Lipofectamine 2000 (ThermoFisher Scientific) following manufacturer's instructions. Lipin1$^{mGFP}$ expression was induced by incubation with 1 µg/ml tetracycline for 24 hours, and cells were fixed 24h later with 4% formaldehyde (Sigma) in 1x PBS.

*MEF lines*

*Tor1a*$^{+/+}$ and *Tor1a*$^{\Delta gag/\Delta gag}$ lines have been previously described (Cascalho et al., 2020). We targeted *Tor1b* using predesigned Alt-R® CRISPR-Cas9 guide RNA (IDT) against exon1 (5'-g GGAACGGCCCCTCAACACGTCGG-3'), and cloned this into pX459 V2.0;pSpCas9(BB)-2A-Hygro (Addgene #62988, puromycin cassette replaced by hygromycin) according to the Zhang lab protocol (Cong et al., 2013). 500ng of *Tor1b*$^{ex1}$-pX459 or empty pX459 were electroporated into 150 000 cells of each MEF line using the Neon™ system (Invitrogen). Cells were placed under hygromycin B selection (120µg/ml, ThermoFisher Scientific) and expanded for 2 weeks. When lines reached confluency, we collected cell pellets, extracted genomic DNA using KAPA Express Extract kit (Sigma), and PCR amplified *Tor1b* exon1 (Table S2) for sequencing. This identified frameshift mutations. We confirmed loss of TorsinB protein by Western blotting. Later, $2 \times 10^6$ cells were electroporated with 2.5 µg of pcDNA5/FRT/TO-Lipin1$^{mGFP}$ or pBudCE4.1-CTDNEP1$^{scarlet}$ using the T20 program of a Nucleofector™ II/2b device (Lonza) and the manufacturer's instructions. Cells were fixed and prepared for imaging 12-24h after transfection.

**Western Blotting**

Cell pellets were lysed in 100µl T-PER (ThermoFisher Scientific) containing 1x protease inhibitor cocktail (Sigma). Protein concentration was measured using the standard BSA method. Next, 30µg of protein was subject to standard SDS-PAGE, transferred to a PVDF membrane, blocked for one hour (5% milk, 0.2% Tween, 1x PBS), and then incubated overnight at 4°C with primary antibodies in blocking buffer. The following day membranes were washed in buffer (0.2% Tween, 1x PBS), and incubated with horseradish peroxidase (HRP) conjugated secondary antibodies diluted in blocking buffer. They then received a final series of washing steps, followed by incubation with West Pico Plus chemiluminescent reagent (ThermoFisher Scientific). Chemiluminescence was detected with an ImageQuant LAS 4000 device.

### Fluorescent labeling

Antibodies and concentrations are provided in Table S3.

*Larval Fat Body*

Larval fillet preparations were fixed at room temperature (RT) in 3.7% formaldehyde (Sigma) in 1x PBS for 20 min, and then washed at room temperature in PBS containing 0.4% Triton X-100 (PBSX). Cell borders and lipid droplets were visualized by sequentially incubating (for 45 min at RT) samples with Phalloidin (red or green; 0.03 mg/ml in 1x PBSX), washing 3 x 10 min in PBSX, incubating with BODIPY 493/503 (1 µg/ml) or HCS LipidTox Neutral Red 577/609 or Deep red 650/670 (1:300) for 30 min at RT and washing 3 x 10 min in PBSX. Immunolabeling was performed as previously described (Soldano et al., 2013). Prior to imaging using a Nikon A1R Eclipse Ti microscope or Zeiss Airyscan, the fat bodies of 4 DO and 5DO animals were removed and mounted in Vectashield with DAPI. 3DO animals were mounted with DAPI as an intact fillet preparation.

*Cultured Cells*

Cells plated on coverslips were fixed with 4% PFA (Sigma) in 1X PBS for 20min at RT. Residual PFA was removed by three 10min washes with 1x PBS. Coverslips were incubated with blocking buffer consisting of 1 X PBS; 0.25% Triton-X100 0nd 10% Normal Donkey Serum (Jackson Immunoresearch) for 30min at RT in a humidified chamber. Then they were placed in a humified chamber and incubated for 16h at 4°C with primary antibody diluted in the same buffer. The following day coverslips received three to five washes of 5 – 10mins, with 1 X PBS, followed by 2 hour incubation at room temperature with secondary antibodies were diluted in blocking buffer. After a final series of three to five washes with 1 X PBS, coverslips were mounted with Vectashield with DAPI (Vector) and imaged.

#### Electron Microscopy

*TEM*

Isolated fat bodies from several animals were fixed in 2.5% glutaraldehyde (Electron Microscopy Services (EMS) #16220), 4% paraformaldehyde (PFA) (EMS #15714), 0.2% picric acid (EMS #19554), 1% sucrose in 0.1 M PB (phosphate buffer, pH 7.4) at 4°C for at least 24 hours, and up to 3-weeks. The tissue was then condensed by embedding in 4% low gelling temperature agarose (Sigma, #A9414) and replaced in fixative at 4°C overnight. Samples were washed three times in 0.1 M SCB (sodium cacodylate trihydrate buffer (pH 7.6); EMS #12300), post-stained with 1% OsO4 (2% Aqueous solution, EMS #19152) and 1.5% potassium ferrocyanide (Sigma #455989) diluted in 0.1 M SCB (pH 7.6) for 1h. Then samples were incubated with 0.2% tannic acid (EMS #2170) and stained with 0.5% uranyl acetate (EMS #22400) in 25% methanol overnight, followed by washing and staining with lead aspartate (EMS #17900) en bloc for 30min at 60°C. Finally, samples were dehydrated through a graded series of ethanol solutions, infiltrated and embedded in Epon (Agar100) and cured at 60°C for 48hrs. Sections (70 nm) were cut with a Dupont diamond knife on a Leica UCT ultra-microtome and collected on copper grids. Sections were imaged with JEOL JEM1400 transmission electron microscope operated at 80kV and equipped with an Olympus SIS Quemesa (11 Mpxl) camera.

*3D-EM*

Larval fillets were fixed in 4% PFA, 0.5% glutaraldehyde in 0.1M PB for 1 hour at room temperature. After samples were rinsed in 0.1M PB, near-infrared branding (NIRB) was performed using a Zeiss LSM 780 inverted confocal microscope equipped with a Mai Tai DeepSee two-photon laser (Spectra-Physics). Branding marks were introduced as a square around nuclei of interest using the two-photon laser at 800nm and 40% maximal power. Two Z-stacks with a 63x objective and a 25x objective 5x zoom were taken of the region of interest after NIRB together with an overview with the 25x objective.

Samples were then trimmed, and post-fixed overnight at 4°C in freshly prepared 4% PFA, 2.5% glutaraldehyde, 0.2% picric acid in 0.1M PB, and stored in this solution until further processing. Next, samples were incubated in 1% $OsO_4$, 1.5% potassium ferrocyanide in ultrapure water for 1 hour at room temperature, followed by 30 minutes in 0.2% tannic acid. Then samples were incubated in 0.5% uranyl acetate in 25% methanol overnight at 4°C. After rinsing with ultrapure water, samples were stained *en bloc* with lead aspartate and dehydrated using ice-cold solutions of increasing ethanol concentration followed by flat embedding in Epon resin. Embedded samples were mounted on aluminum pin stubs (Gatan) with conductive epoxy glue (Circuit Works) and inserted in a Zeiss Sigma Variable pressure SBF-SEM with 3View technology (Gatan) to approach the region of interest. Once nuclei of interest were located, based on the branding marks and fat body morphology, the pins were inserted in FIB-SEM. For FIB-SEM imaging, samples mounted on pins were coated with ~20nm of platinum in a sputtercoater (Quorum Q150T ES). FIB-SEM imaging was performed using a Zeiss Crossbeam 540 system with Atlas5 software. The Focused Ion Beam (FIB) was set to remove 5nm sections by propelling Gallium ions at the surface. The Atlas software was set to image an area of 10 x 5 μm area at 5nm pixels (5μs dwell time &

line averaging 4) using an ESB (back-scattered electron) detector with the electron beam at 1.5 kV and 1 nA.

FIB-SEM data was first denoised using Fiji and the Tikhonov algorithm using following parameters; lambda =2, iterations=50 and sigma=1.058. Next, data was segmented using Microscopy Image Browser (MIB) software and 3D-modeling was performed using Amira software (FEI/Thermo Fisher Scientific, France).

*CLEM*

Larval fillets were fixed in 4% PFA, 0.5% glutaraldehyde in 0.1M PB for 30 minutes at room temperature. After samples were rinsed in 0.1M PB and a 1.35mm disc was cut out of larval FB and submerged in 20% BSA in DMEM medium with 10% fetal calf serum (FCS). Next tissue discs were loaded in membrane carriers (0.1mm think, Leica-microsystems), frozen with a high-pressure freezer (Leica EMPACT2) and vitrified at 2050 bar. The quick freeze substitution protocol was started by transferring the membrane carriers to cryotubes (72.694.005, Sarstedt, Germany) with freeze-substitution medium with 0.2% uranyl acetate ((#02624-AB, SPI) in acetone (#1002990500, Merck) and 5% MilliQ water at -180°C. Once the cryotubes reached -80°C the cryotube-holder was placed on its side and agitated. At -50°C, the carriers were transferred to the pre-cooled -50°C Leica AFS2 apparatus, washed in 100% acetone, and after a final rinsing in ethanol, infiltrated with lowicryl HM20 (#02628-AB, SPI), and finally polymerized at -50°C with UV. 200nm sections were cut on a Leica Ultracut S ultramicrotome and collected on microscopic slides. Immediately followed by 70nm sections collected on slot grids (#01805-F, ted pella, USA). The fluorescence image of the last 200nm section was overlaid with EM images of the first 70 nm section using GNU Image Manipulation Program (GIMP).

**Image Analysis**

All analyses were performed using FIJI, by a researcher blind to genotype, and at least three animals and/ or independently prepared cell preparations were examined for each genotype/ condition.

*Cell size and Lipid Droplet density*

The fat body labeled with DAPI, Phalloidin (red or green), BODIPY or LipidTox was imaged at 40x or 60x magnification using a Nikon A1R Eclipse Ti microscope. Cell size was measured by manually outlining individual cells (defined by phalloidin) with the Freehand selection tool, followed by the set measurements > area function. Measurements were only made from cells if the optical section bisected the nucleus. LD size was measured using an optimized macro to quantify single LD particles (analyze particles size= 5-infinity) after smoothening and applying a threshold that was manually determined to highlight most LD in an image. LD density was quantified by manually outlining an individual cell as a ROI, applying a threshold to highlight all green / red pixels inside this region, and measuring the area occupied by fluorescence compared to the total area of the cell.

*Lipin localization*

Fly fat bodies expressing Lipin$^{V5\text{-}His}$ and labeled with anti-V5 and DAPI were imaged at 63X magnification with a Zeiss AiryScan. Nuclei were identified in the DAPI channel, and a ROI was drawn in the interior of every nucleus present in an image. The intensity of anti-V5 staining in the ROI was then measured. Background anti-V5 staining was determined by measuring anti-V5 staining intensity in at least 10 large Lipid Droplets in each image.

HEK293T-Lipin1$^{mGFP}$ cells transfected with pBudCE4.1 plasmids and labeled with anti-myc and anti-V5 were imaged at 60x on a C2 Nikon confocal microscope. MEF lines expressing Lipin1$^{mGFP}$ were imaged at 60x with oil on a A1R1 Nikon confocal microscope. GFP localization was scored as primarily nuclear or primarily cytosolic by an observer blind to condition. This was restricted to anti-myc/V5 positive HEK cells. Cells with obvious abnormalities, including multi-nucleated cells, were not considered. More than 50 transfected cells were examined per condition

*Nucleoporin localization*

Fat body tissue labeled with mAb414 and DAPI was examined with a Zeiss Airyscan and 63x magnification. Individual nuclei were scored for whether mAb414 labeling was exclusively localized to the NE (strong NE staining), showed some NE labeling against a background of cytosolic/ nuclear signal, or had no obvious NE labeling.

*Nuclear membrane ultrastructure*

We defined NE structure as “normal” if the INM and ONM were adjacent, roughly parallel, and no additional membrane was present within the NE lumen. An observer determined the length of NE that fulfilled these criteria, for individual nuclei imaged with TEM at 2500x and 10000x magnification. The entire NE length present in an image was then measured to determine the “percentage of normal NE”. The distance between INM and ONM adjacent to pores/ channels was calculated from the 3D reconstruction of FIB-SEM images. The density of mature nuclear pores and attached blebs (per 10 $\mu$m) were calculated by counting their number in each nuclear profile present in a TEM image at 10000x magnification, as well as measuring the length of the NE that was examined.

### TAG Measurement

TAG was measured as previously described (Palanker et al., 2009). 3DO larvae were homogenized in PBS + 0.05% Tween 20 (PBST) and immediately incubated at 70°C for 10 min. 20µl of the heat-treated homogenates were incubated with 20µL of either PBST or "Triglyceride Reagent" (Sigma; T2449) for 45 min at 37°C, after which the samples were centrifuged at maximum speed for 3 min. Next, 30 µl of the supernatant was transferred to a 96-well plate and incubated with 100 µl of "Free Glycerol Reagent" (Sigma; F6248) for 5 min at 37°C. Finally, absorbance at 560 nm was measured using a Victor3 plate reader spectrophotometer (Perkin-Elmer). TAG amounts were determined by subtracting the amount of free glycerol in the PBST-treated sample from the total glycerol present in the sample treated with Triglyceride Reagent. Data is represented relative to control.

### Statistical analyses

Statistical analyses were performed using GraphPad Prism 8.2.1 software. The criteria for significance are: ns (not significant), * $p < 0.05$, ** $p < 0.01$, *** $p < 0.001$, *** $p < 0.0001$. Differences between groups were assessed by One-way ANOVA, and Bonferroni's post hoc test, unless otherwise stated in figure legend.
