## Supplementary material for "The Torsin/ NEP1R1-CTDNEP1/ Lipin axis regulates nuclear envelope lipid metabolism for nuclear pore complex insertion": Fig. S3

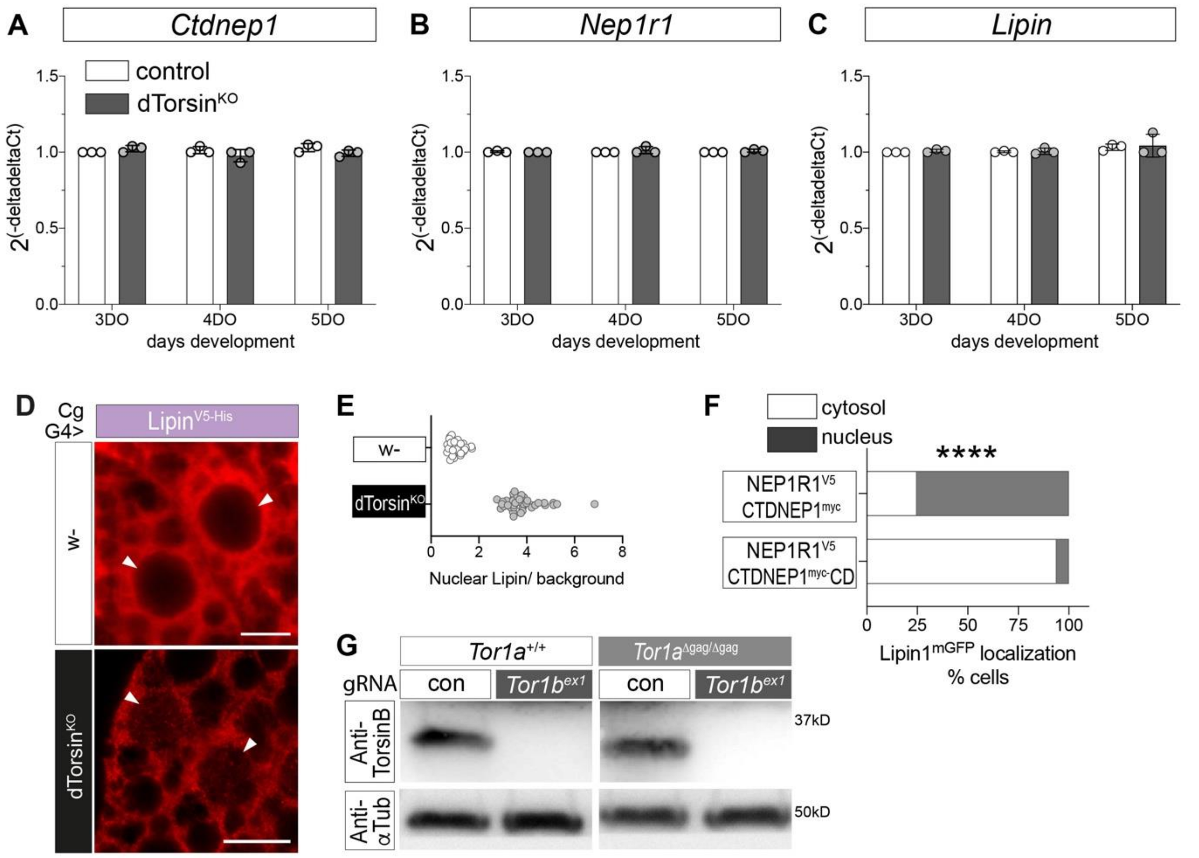


**Figure S3**

**Related to Figure 6: Torsins control the ER/NE localization of CTDNEP1, and the cytosolic/nuclear localization of Lipin**

A - C) qRT-PCR of cDNA prepared from fat bodies of 3DO, 4DO and 5DO control and *dTorsin*$^{KO}$ larvae. Bars show group mean, points show individual values from *n = 3* independently run qRT-PCR.

D) Anti-V5 labeling of 4DO wild-type (w-) or *dTorsin*$^{KO}$ fat body cells expressing Lipin$^{V5-His}$. Arrowheads indicate nuclei. Scale bars, 10μm.

E) Ratio in individual cells between nuclear V5 signal and LD V5 signal (background).

F) Percentage of HEK293-Lipin1$^{mGFP}$ cells with primarily cytosolic GFP signal (white bars) or nuclear GFP signal (dark bars) after transfection with the dual promoter pBudCE4.1 containing NEP1R1$^{V5}$ and either CTDNEP1$^{myc}$ or CTDNEP1$^{myc}$-CD. **** indicates significant difference (Fisher's exact test).

G) Anti-TorsinB and anti-$\alpha$Tubulin (Tub; loading control) Western Blotting of lysates from wild-type Mouse Embryonic Fibroblasts (MEFs) and *Tor1a*$^{\Delta gag/\Delta gag}$ MEFs after electroporation and selection for CRISPR plasmids. *Tor1a*$^{\Delta gag/\Delta gag}$ is a strong loss-of-function allele (Goodchild et al., 2005) and we refer to these as *Tor1a*$^{KO}$.
