## Supplementary material for "The Torsin/ NEP1R1-CTDNEP1/ Lipin axis regulates nuclear envelope lipid metabolism for nuclear pore complex insertion": Fig. S4

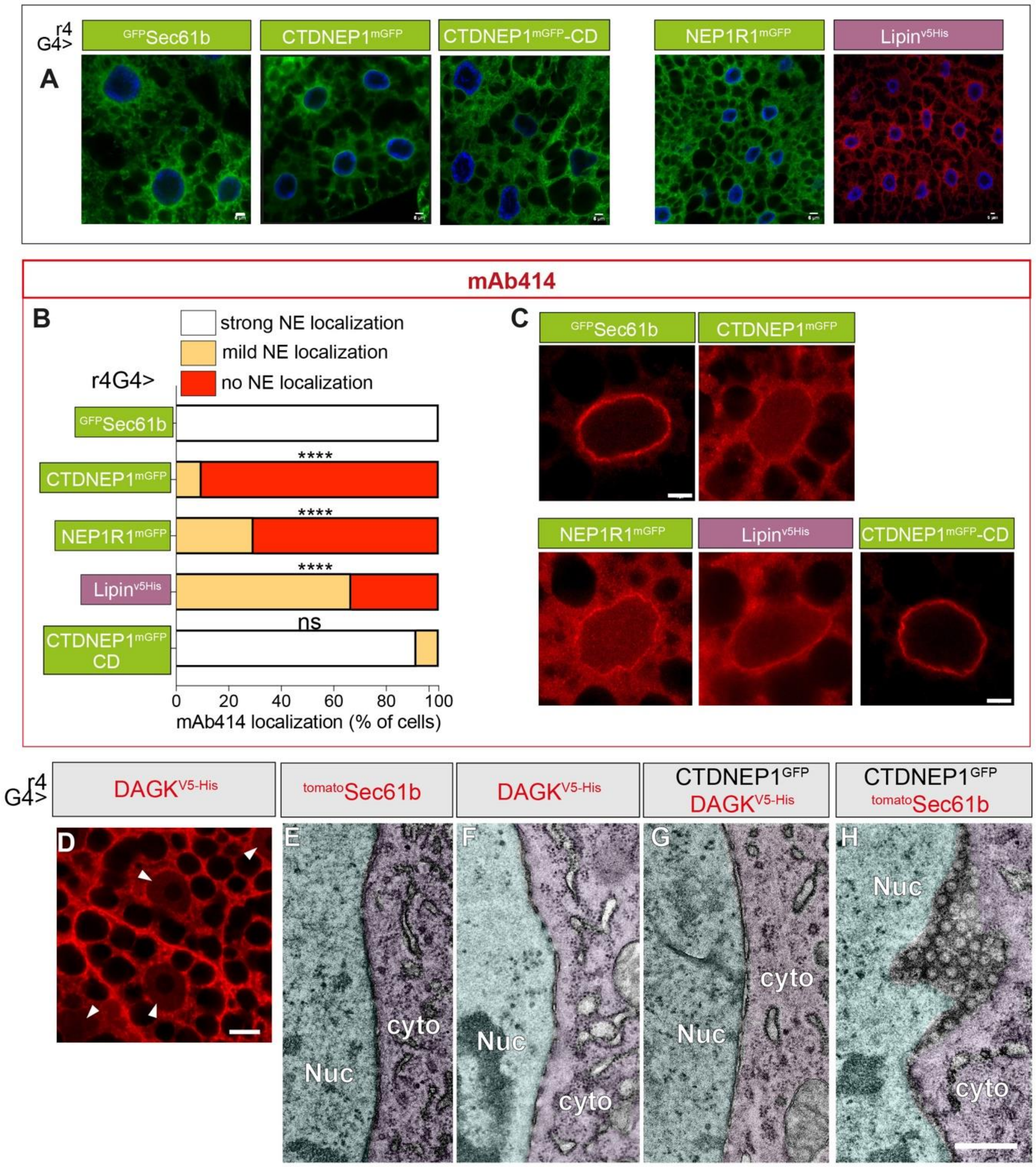


**Figure S4**

**Related to Figure 7: CTDNEP1 phosphatase activity inhibits NPC maturation by dysregulating lipid metabolism**

A) Representative confocal images of fat body cells expressing $^{mGFP}$Sec61b (control), CTDNEP1$^{mGFP}$, CTDNEP1$^{mGFP}$-CD, NEP1R1$^{mGFP}$ or Lipin$^{V5\text{-}His}$. Scale bar, 5μm.

B) Quantification of mAb414 localization in 5DO fat body cells expressing each transgene. White bars show the percentage of cells with intense NE-localized mAb414 signal, orange shows the percentage with low but visible NE mAb414 signal, and red shows the percentage of cells where NE mAb414 signal was absent. > 50 cells from three separate larvae were examined for each condition. **** indicates a significant difference in the number of cells with strong NE labeling compared to $^{mGFP}$Sec61b (control). Chi square test ($p < 0.0001$), followed by individual post hoc Chi square tests.

C) Representative confocal imaging of mAb414 localization in fat body cell nuclei expressing $^{mGFP}$Sec61b (control), CTDNEP1$^{mGFP}$, NEP1R1$^{mGFP}$, Lipin$^{V5\text{-}His}$ or CTDNEP1$^{mGFP}$-CD. Scale bar, 5µm.

D) Confocal imaging detects V5 signal in the cytosol and nucleus (arrowheads) of fat body cell nuclei expressing UAS-DAGK$^{V5\text{-}His}$ with r4-Gal4. Scale bar, 10µm.

E – H) NE ultrastructure in 5DO fat body cells expressing UAS- transgenes with r4-Gal4. Scale bar, 500nm.
