## Supplementary material for "The Torsin/ NEP1R1-CTDNEP1/ Lipin axis regulates nuclear envelope lipid metabolism for nuclear pore complex insertion": Fig. S2

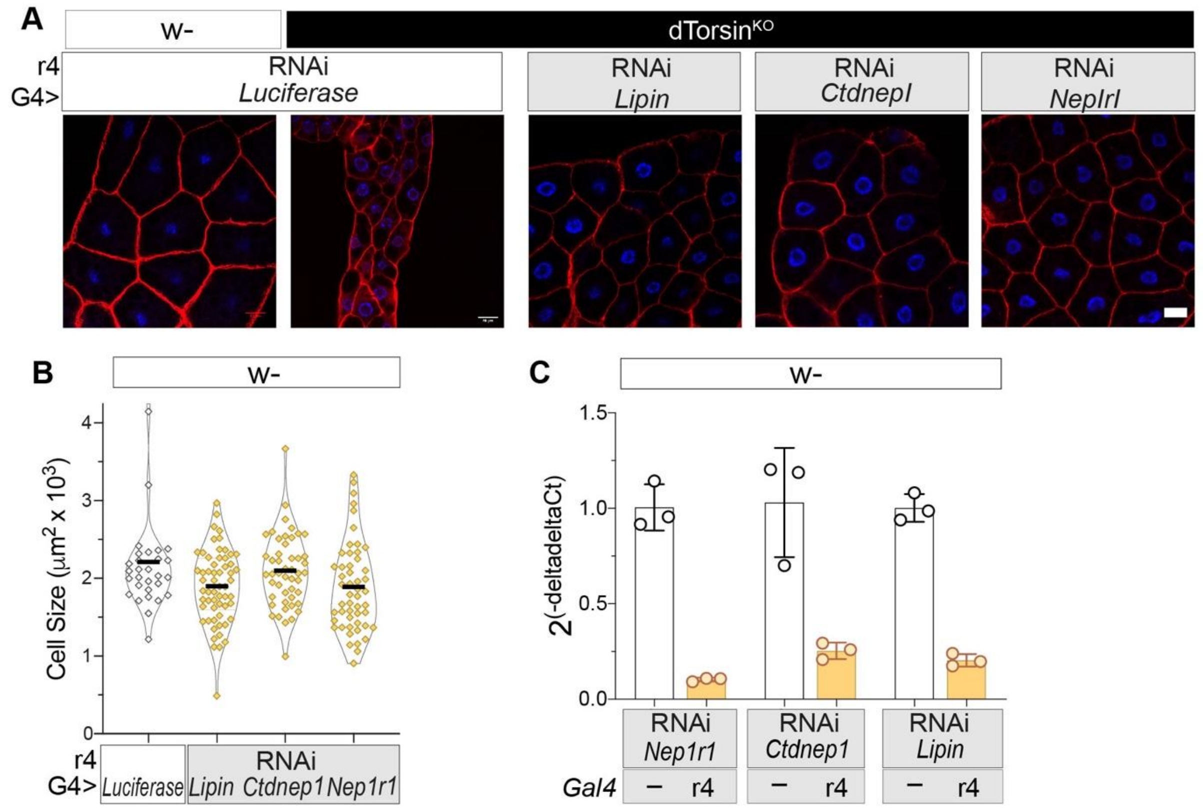


**Figure S2**

**Related to Figure 3: *Nepr1/2-Ctdnep1* RNAi suppresses cell growth, organelle volume, and LD defects of *dTorsin*$^{KO}$ larvae**

A) Confocal images of Phalloidin and DAPI stained wild-type (w-) or *dTorsin*$^{KO}$ fat body cells expressing UAS-RNAi transgenes with r4-Gal4. Scale bar, 20μm.

B) Points show values from individual fat body cells expressing the indicated UAS-RNAi transgene with r4-Gal4. Bars show the group mean.

C) qRT-PCR of cDNA prepared from fat bodies of 5DO larvae with UAS-*Lipin* RNAi, UAS-*Ctdnep1* RNAi, and UAS-*Nep1r1* RNAi, with and without the r4-Gal4 transgene. Bars show group mean, points show individual values. *n = 3* independent qRT-PCR.
