## Supplementary material for "The Torsin/ NEP1R1-CTDNEP1/ Lipin axis regulates nuclear envelope lipid metabolism for nuclear pore complex insertion": Fig. S1

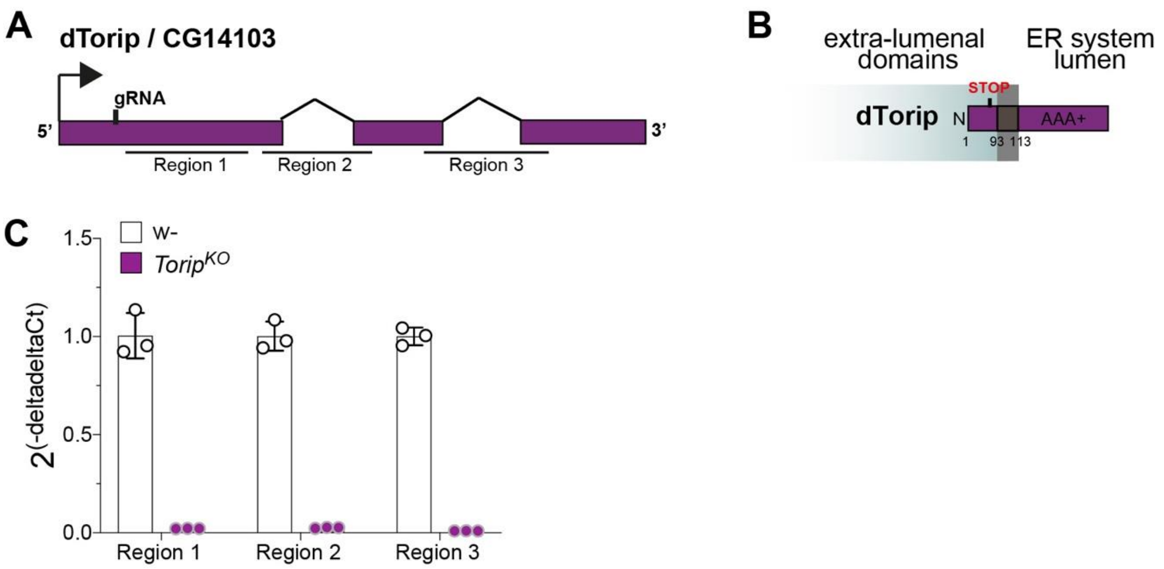


**Figure S1: Characterization of the *Torip* Δ11 allele**

**Relates to Figure 2: *dTorsin* and *Torip/LAP1* independently regulate lipid metabolism at the nuclear envelope**

A) *Torip* gene structure and site of guide RNA used for CRISPR gene editing.

B) Torip protein structure, highlighting the premature stop codon (position 71) generated by the frameshift Δ11 mutation.

C) qRT-PCR from 5DO 3rd instar control or *Torip*$^{\Delta 11/\Delta 11}$ larvae with primers that amplify 3 regions of the *Torip* cDNA. Graph shows mean +/- SD of n = 3 replicates.
